# Insulin signaling and extended longevity in ants

**DOI:** 10.1101/2022.06.25.497611

**Authors:** Hua Yan, Comzit Opachaloemphan, Francisco Carmona-Aldana, Giacomo Mancini, Jakub Mlejnek, Nicolas Descostes, Bogdan Sieriebriennikov, Alexandra Leibholz, Xiaofan Zhou, Long Ding, Maria Traficante, Claude Desplan, Danny Reinberg

**Affiliations:** Department of Biochemistry and Molecular Pharmacology, New York University School of Medicine, New York, NY 10016, USA; Howard Hughes Medical Institute, New York University School of Medicine, New York, NY 10016, USA; Department of Biology, Center for Smell and Taste, University of Florida, Gainesville, FL 32611, USA; Department of Biology, New York University, New York, NY 10003, USA; Guangdong Laboratory for Lingnan Modern Agriculture, Guangdong Province Key Laboratory of Microbial Signals and Disease Control, Integrative Microbiology Research Centre, South China Agricultural University, Guangzhou 510642, China

## Abstract

In most organisms, the cost of reproduction is a shorter lifespan. However, the reproductive caste in eusocial insect species (queen) exhibits an extraordinarily longer lifespan than non-reproductive castes (workers) despite having a similar genome, thus contradicting the aging dogma. In the absence of the queen, *Harpegnathos saltator* ants can undergo a caste switch from workers to reproductive pseudo-queens (gamergates). Gamergates exhibit a dramatically prolonged lifespan. When placed in the presence of a reproductive, they revert to worker status and their lifespan is then shortened accordingly.

To understand this unique relationship between reproduction and longevity, we analyzed tissue-specific gene expression between castes. Insulin is upregulated in the gamergate brain that leads to oogenesis, but surprisingly correlates with extended longevity. This correlates with increased lipid synthesis and elevated production of vitellogenin in the fat body, which are both transported to the egg. We show that the production of vitellogenin in the fat body is due to the systemic activation of the MAPK branch of the insulin/IGF signaling (IIS)-pathway. In contrast, reduced expression of insulin receptors in the fat body of gamergates and the production in their developing ovary of an anti-insulin (Imp-L2) lead to the downregulation of the AKT/FOXO branch of the IIS-signaling pathway in the fat body, and to the dramatically extended longevity. This reveals a dual-pathway mechanism that reconciles increased longevity in the context of active reproduction in eusocial insects.

**One Sentence Summary:** Insulin-dependent reproduction in ants correlates with extended longevity through insulin inhibition by anti-insulin Imp-L2.

## Main text

Differences in lifespan within a species offer a comparative paradigm to investigate the underlying regulatory processes involved in aging. Reproduction has an important influence on longevity: genes that favor reproductive fitness normally shorten lifespan (*1, 2*) as animals allocate nutritional and metabolic resources for reproduction at the cost of longevity (*3*). This idea is corroborated by the adaptive responses to dietary restriction in different species, which include reduced reproductive capability and prolonged lifespan (*4, 5*). The functional anticorrelation between reproduction and longevity involves the insulin/IGF signaling pathway (IIS) as increased IIS activity required for reproduction leads to shorter lifespan in most animals (*1, 6-9*).

Extensive studies performed in *C. elegans, Drosophila*, mouse and other model organisms have analyzed the effects of signaling pathways, notably the IIS pathway, in regulating longevity (*10-13*). In insects, the brain, fat body (the metabolic organ of insects that is equivalent to the vertebrate liver and adipose tissue), and ovary are the primary tissues regulating longevity and reproduction (*14, 15*). Ablation at the larval stages of the insulin producing cells (IPCs) in the *Drosophila* brain causes lower female fecundity, elevated lipid storage lipids, and extended lifespan (*16*). The fat body mainly contains adipocytes and oenocytes (hepatocyte-like cells), which play essential roles in energy storage and utilization (*17, 18*), pheromone synthesis, reproduction and longevity (*15, 17, 19-21*). Repression of IIS through upregulation of FOXO, the negative effector of IIS, in the *Drosophila* fat body or adipose tissue in mice extends their lifespan, but reduces fecundity (*10, 21-24*). The ovary also plays a role in regulating lifespan: removal of germline cells extends lifespan in *Drosophila* and *C. elegans (25-27)*. However, eusocial insects, like ants, bees, wasps and termites, present a striking paradox to the coupling of reproduction and longevity: the complete reproductive function of a colony depends on one or very few female individuals (queens) that exhibit a dramatically increased longevity (up to 30x) compared to their non-reproductive female nestmates (workers) (*28-31*), despite sharing a similar genome. Yet it is not clear how this reproduction-associated longevity is regulated at the cellular and molecular levels. This paradox implies that reproduction and lifespan are uncoupled in eusocial insects. Alternatively, it suggests an intriguing possibility that a protective mechanism allows for prolonged lifespan in the reproductive caste.

### Reversible longevity in the reproductive caste

To understand the mechanisms that give rise to this paradox, we used the ant *Harpegnathos saltator* that exhibits a particular trait: when the queen in a colony dies, non-reproductive workers start dueling with each other. The winners gradually transition to so-called gamergates (pseudo-queens) that continue dueling, develop queen-like behavior, begin laying eggs and exhibit a dramatic lifespan extension (Fig. 1A) (*32-35*). Gamergates can also be reverted back to workers (‘revertants’) when they are placed in a colony with an established reproductive caste, leading to a shortened lifespan (*36*). The caste switching plasticity of this species offered a prime opportunity to investigate the coupling between reproduction and lifespan. While the median lifespan of workers is 217 days (Fig. 1B and S1A), the median lifespan of gamergates was not reached in our observation period of ∼600 days, which is in accordance with other work showing that it is ∼1,100 days (*29*). The median lifespan of revertants was 188 days, which is comparable to 184 days found in their worker nestmates (Fig. 1B and S1A). This is shorter than the 217 days of regular workers, presumably because the policing of gamergates by workers during the reversion induces colony stress.

**Fig. 1.**
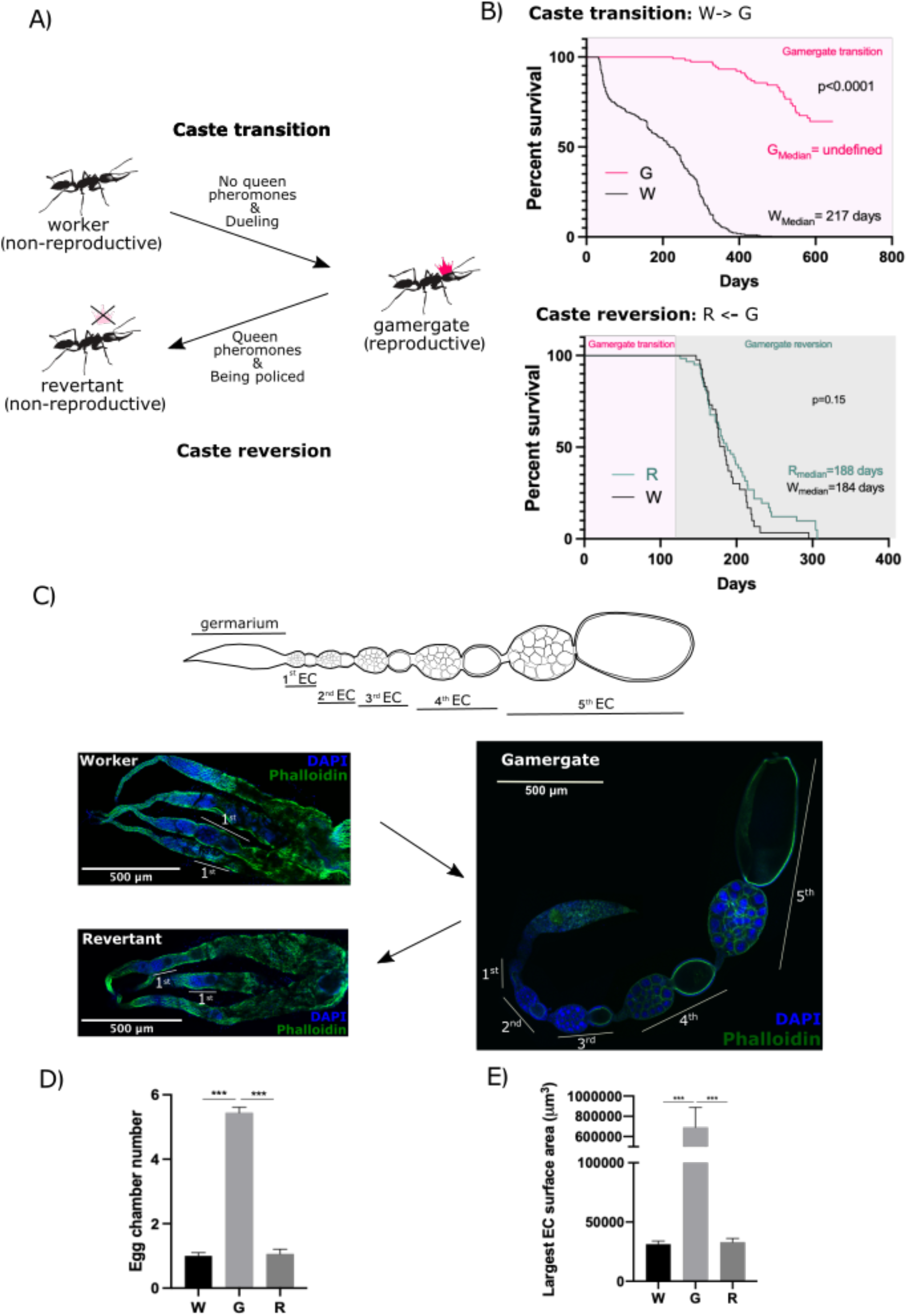
Phenotypic plasticity in the ant *Harpegnathos saltator*. (**A)** *Harpegnathos* female workers retain reproductive potential, which is suppressed by queen pheromone. Removal of queen pheromone drives some non-reproductive workers to start dueling and become reproductive pseudo-queens, a.k.a. gamergates (caste transition). The gamergates can transition anew to non-reproductive revertants in the presence of queen pheromones (policing and caste reversion). (**B**) Lifespan of ants during the caste transition and reversion. Upper panel: survival curves of non-reproductive workers (W, black, n=291) vs. reproductive gamergates (G, pink, n=40) during the W-to-G transition. Lower Panel: survival curves of revertants (R, green, n=47) vs. workers (W, black, n=36) during the G-to-W reversion. Gamergates derived from 3 months of transition (pink box) were subsequently subjected to reversion (green box). p values from Log-rank (Mantel-Cox) test are indicated in the plots. (**C**) Ovary development during the caste transition and reversion. Upper panel: schematic of an ovariole within a gamergate ovary comprising a germarium and different stages of developing egg chambers (ECs). The 6^th^ EC, which only contains a large oocyte without nurse cells, is not shown. Lower panel: immunofluorescence (IF) staining of ovaries of worker (top-left) and revertant (bottom-left) and a single ovariole of a gamergate (right) with Phalloidin (green) and DAPI (blue). The numbers of developing ECs are indicated in the gamergate panel. (**D-E**) Quantifications of the average number of developing ECs per ovariole (**D**) and the surface area of the largest EC (**E**) in each ovariole of worker (W), gamergate (G) and revertant (R). p-values are from Kruskal-Wallis test and multiple comparisons (p<0.001***, n=5). Bars and error bars represent mean ± SEM.

To determine the reproductive potential of the castes, we dissected the gaster (abdomen) and inspected the ovaries. Fertile gamergates had eight ovarioles (four on each side) that contained chains of 4-6 egg chambers including some with fully developed oocytes (stage 6) (Fig. 1C, D). In contrast, workers had ovaries with very small, partially developed ovarioles that each contained the germarium where stem cells are located, with none to two (0-2) early-stage egg chambers (stages 1 or 2) and no fully developed oocytes. Thus, ovary growth is blocked at the early stage of egg formation in workers, but can be reactivated in gamergates (Fig. 1C, D). The ovaries of revertants also had 0-2 immature egg chambers per ovariole (Fig. 1, C-E), which correlated with their loss of reproduction (*36*). In summary, reproduction and longevity are positively coupled in *Harpegnathos*: as animals switch caste and start producing many eggs, their lifespan increases, and *vice versa*. We thus sought to address how lifespan can be increased during active reproduction in this species, which is in very sharp contrast with most model organisms.

### Upregulation of insulin in the reproductive caste

We performed bulk RNA-sequencing from tissues that are important for reproduction and metabolism (brain, ovary and fat body) from workers, gamergates and revertants. To validate the accuracy of our transcriptomes, we confirmed the differential expression profile of already characterized genes, such as the gene encoding the neuropeptide Corazonin (Crz) that is highly expressed in the worker brain, and the egg yolk precursor *Vitellogenin* (*Vg*) gene that is increased in the gamergate fat body (*37*) (Fig. 2A, B, S1B).

**Fig. 2.**
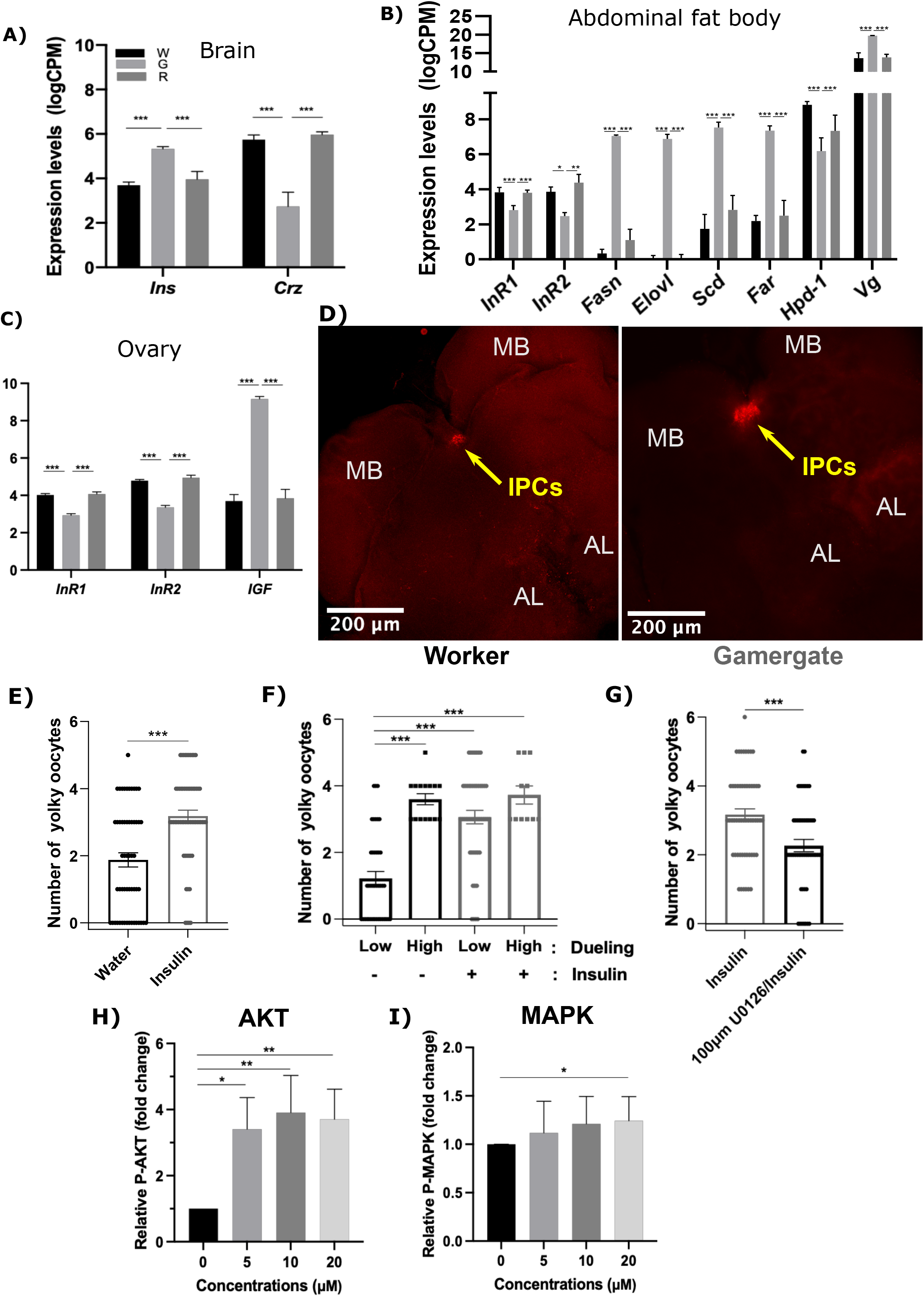
Ins-related gene expression in different castes and its induction of oogenesis via IIS-MAPK pathways. (**A-C**) RNA-seq levels of differentially expressed genes (DEGs) in the central brain (**A**), abdominal fat body (**B**), and ovary (**C**) in workers (W, black), gamergates (G, grey) and revertants (R, dark grey). Data are from four biological replicates per caste. p values from EdgeR are indicated. *Hpd-1*: *4-hydroxyphenylpyruvate dioxygenase, Fasn*: *fatty acid synthase, Elovl*: *fatty acid elongase, Scd*: *desaturase, Far*: *fatty acyl-CoA reductase*. (**D**) Localization of *Ins* mRNA in the worker (left panel) and gamergate (right panel) brains by *in situ* hybridization (ISH). *Ins* mRNA is located in the insulin producing cells (IPCs, indicated by yellow arrows) located between mushroom bodies (MB). AL: antennal lobe. (**E-G**) Yolky oocyte production in control vs. Ins-injected ants (**E**), in low-vs. high-dueling ants (**F**), and in Ins-injected vs. Ins and U0126 co-injected ants (**G**). In (**E**) and (**G**), ovary development is represented by the number of yolky oocytes per individual in all workers regardless of their dueling activity. Bars and error bars represent mean ± SEM (p<0.05*, 0.01**, 0.001***). (**H-I**) Quantification of fold changes in relative phosphorylated AKT levels (P-AKT/T-AKT, **H**) and phosphorylated MAPK levels (P-MAPK/T-MAPK, **I**) in fat body as a function of different concentrations of insulin treatment. Relative levels of AKT and MAPK phosphorylation were normalized to the control fat body. n=8, p values are from Kruskal-Wallis test with multiple comparisons. Bars and error bars represent mean ± SEM.

The IIS pathway has a central role in metabolism and in particular, in female reproduction. Importantly, mutants in the IIS pathway exhibit extended lifespan in most species studied but also reduced reproduction (*10, 38*). To better understand insulin signaling in eusocial insects, we performed a phylogenetic analysis of genomic and transcriptomic data of 36 Hymenopteran species (ants, bees and wasps). We found that in almost all species, including *Harpegnathos*, but not in the ant *Camponotus floridanus*, there are 2 insulin-like peptides (ILPs): an insulin homolog (Ins) and an insulin-like growth factor homolog (IGF) (Fig. S2A, B). Ins contains only A and B chains, while IGF has A, B and retains the C chains. Both ILPs can form 3 disulfide bonds, a structure essential for their interactions with receptors (Fig. S2C, D). We identified several differentially expressed genes related to the IIS pathway in *Harpegnathos*: *Ins* was upregulated in the gamergate brain, while *IGF* was upregulated in the mature gamergate ovary (Fig. 2A, C). In contrast, the two genes encoding an Insulin Receptor (InR) were downregulated in the gamergate fat body and ovary (Fig. 2B, C), but were unchanged in the brain. We generated mRNA probes and antibodies against *Harpegnathos* Ins and IGF and examined their distribution by *in situ* hybridization and immunofluorescence staining. As in *Drosophila*, the main source of Ins in the brain are the insulin producing cells (IPCs) (*39*) (Fig. 2D and S3A). *Ins* mRNA was expressed at much higher levels in the cell body of IPCs in gamergates than in workers (Fig. 2D). Ins protein accumulated in both the cell body and the axons of the IPCs (Fig. S3A). Increased *IGF* mRNA and protein were mainly detected in nurse cells and the follicle cells of the gamergate ovary (Fig. S3B). This increase in ILPs in gamergates is consistent with the high metabolic requirement for egg production. However, high IIS activity should lead to decreased lifespan.

### Altered metabolism in gamergates

As IIS regulates metabolism, we next analyzed metabolic changes in gamergates. The gamergate fat body exhibited increased expression of genes related to lipid metabolism and modifying enzymes, such as *fatty acid synthase* (*Fasn*), *fatty acid elongase (Elovl), desaturase* (*Scd*) and *fatty acyl-CoA reductase* (*Far)*, suggesting an active synthesis of lipids either for the production of the egg yolk, or of cuticular hydrocarbon (CHC) that comprise long-chain hydrocarbons and constitute the queen pheromones (*35, 36, 40-45*) (Fig. 2B). Thus, the fat body of gamergates exhibits transcriptional signatures of increased lipid production. The fat body was large and white-colored in the abdomen of workers, while it was greatly reduced in size and had a yellowish color in the gamergate abdomen that was mainly occupied by the developed ovarioles (Fig. S4A). The dissected worker fat body floated in saline solution (1xPBS), but the gamergate fat body sank (Fig. S4B), reflective of a lower lipid content in the latter case. Indeed, we measured the lipid contents [tri- and di-acylglycerol (TAG and DAG, respectively)] using an enzymatic conversion of glycerides into glycerol. Lipids were significantly reduced in the gamergate fat body as compared to workers (Fig. S4C). However, in line with the upregulated *Fasn* expression in the fat body, circulating lipids were increased in the hemolymph of gamergates, pointing to lower lipid storage in favor of high lipid mobilization in the reproductive gamergates, likely for their utilization by the ovary for oogenesis (Fig. S4D, E). Moreover, Nile Red staining in the fat body revealed that lipid droplets were abundant in the enlarged adipocytes of workers, but rarely detected in the smaller adipocytes of gamergates. Unlike in workers, lipids were mainly found in gamergate oenocytes (Fig. S4, F-H). This result is consistent with the increased production of queen pheromone in gamergates as oenocytes are the main source for CHC pheromone synthesis in the abdomen (*17, 43, 46*). In addition to lipids, we also measured the carbohydrate levels in dissected fat body tissue by treating with the enzyme trehalase that converts trehalose into glucose, followed by a hexokinase assay to determine glucose levels (see Methods). Glucose and trehalose levels were decreased in the fat body after the gamergate transition, while their circulating levels in the hemolymph were unchanged (Fig. S4I, J). Thus, the gamergate fat body has reduced carbohydrate storage, which is opposite to expected given the increased production of ILPs in the brain and ovary that should increase IIS and carbohydrate storage. It appears that the ILPs are not acting on the fat body and are less active in gamergates than in workers despite their increased production. These altered carbohydrate and lipid contents in the gamergate fat body may reveal clues to the fecundity-induced longevity in *Harpegnathos*.

### Ins does not induce dueling, but promotes oogenesis in workers

Ins and IGF activate ovary growth in multiple invertebrate and vertebrate species, and Ins induces reproduction in clonal raider ants (*47*). To test the role of IIS in *H. saltator* reproduction, we synthesized its Ins. We injected the Ins peptide into the gaster of ∼2 week old workers in a queen-less colony wherein workers start dueling to initiate the transition to gamergate (*32-35*): two days after the setup of 3 independent colonies, half of the members (20 individuals) of each colony were injected with the Ins peptide while the others were injected with a control solution. We scored the dueling behavior 5 days after dueling initiation and measured the development of the ovary. Ins injection did not induce dueling behavior in workers (27% of workers were dueling in control while 19% of workers dueled in the Ins group, n=3 colonies, p=0.75, Fig. S5A). However, we did observe significant development of the ovarioles in injected individuals with an average of 3.2 egg chambers as compared to 1.9 in control ants (Fig. 2E). Normally, in this experimental context, dueling activity positively correlates with the probability of transitioning to a gamergate, while low-dueling workers do not normally have developed ovaries. However, we found that the number of yolky oocytes was increased by ∼2.3x in the low-dueling workers after injection with Ins peptide (3.0 vs. 1.3, p<0.001), which was mainly ineffectual in the case of high-dueling individuals (3.8 vs. 3.6, Fig. 2F). As expected, dueling and oocyte numbers were positively correlated in control workers (R^2^=0.54), but were not correlated in Ins-injected workers (R^2^=0.02, Fig. S5B). These findings suggest that although Ins could not induce dueling in workers, it was still able to promote oogenesis independently of the dueling status (Fig. 2, E-G and S5A, B). We also injected Ins into the workers in the colonies with established reproductives wherein any worker that attempts to become gamergate is subject to policing due to altered CHC profile (*36, 43*). Interestingly, Ins-injected ants were not policed while they developed ovarioles (Fig. S5C), suggesting that pheromone production is not directly controlled by Ins and ovarian development. In clonal raider ants, Ins injections also promote egg production although in a different context (*47*).

### Ins can activate AKT and MAPK

Thus far, our experiments showed that high Ins in gamergates promotes oogenesis, but does not induce carbohydrate storage and instead, promotes fat body production of vitellogenin and of lipids that are likely transported to the hemolymph for egg formation. This suggested that only part of the IIS pathway is functioning as expected. Thus, we next tested whether Ins might differentially activate distinct branches of the IIS pathway. ILPs can induce AKT phosphorylation that prevents nuclear localization of the Forkhead box O (FOXO) transcription factor. Thus, nuclear FOXO is negatively regulated by IIS and promotes longevity. ILPs can also induce phosphorylation of MAPK/ERK to increase cell proliferation (*48-51*). We therefore treated *ex vivo* dissected worker fat bodies with the synthetic Ins peptide. Ins was able to strongly activate phosphorylation of AKT (∼3.5x increase after Ins treatment, p<0.01, n=8, Fig. 2H and S5D). We also tested the effect of Ins on the MAPK pathway using phosphorylated MAPK/ERK as a readout. The MAPK pathway was mildly activated in the Ins-treated fat body of workers (∼1.3x increase after treatment, p<0.05, n=8, Fig. 2I and S5D). Although MAPK can be activated by Ins, the Epidermal Growth Factor receptor (Egfr) is the most common activator of this pathway, although other Receptor Tyrosine Kinases also induce MAPK (*52, 53*). However, our transcriptome indicated that *Egfr* and its ligands, *Spitz* and *Vein*, were downregulated in the gamergate fat body while *Vein* was also repressed in the ovary (Fig. S1B), suggesting that the EGF pathway does not play a major role in activating MAPK in the fat body. However, other unidentified ligands besides Ins might also contribute to MAPK activation.

### AKT is downregulated in the gamergate fat body, but not in the ovary

As Ins activates AKT and MAPK (Fig. 2H, I, and S2D), we expected to see global upregulation of both IIS pathways *in vivo*. To this end, we dissected multiple tissues, including the fat body and ovary, and analyzed the activity of MAPK using its phosphorylated state as a readout (Fig. S2D). Consistent with the elevated level of Ins in gamergates, MAPK activity was increased in the gamergate fat body (approximately 4x increase, n=6, p<0.001, Fig. 3A, B) and in the ovary, including the germarium and the early- and late-stage egg chambers, and in the malpighian vesicle (MV), the tissue equivalent to the vertebrate kidney (Fig. 3A and S5E). However, MAPK activity was unchanged in the brain and the post pharyngeal gland (PPG), a tissue likely involved in the synthesis and storage of cuticular hydrocarbon (CHC) pheromones (Fig. S5E) (*54, 55*).

**Fig. 3.**
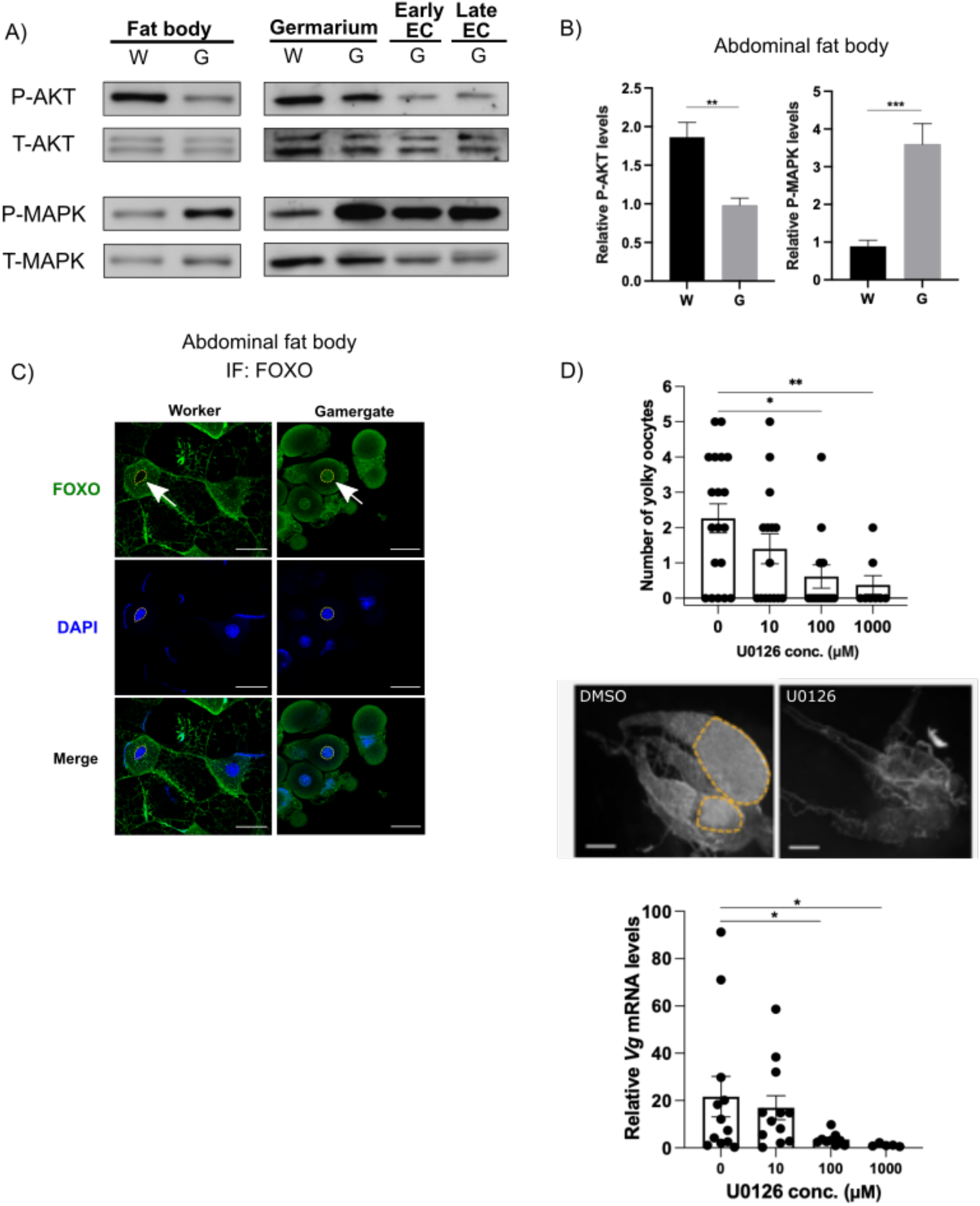
Activity of the IIS-AKT/-MAPK pathways in different tissues and the requirement of MAPK activity for reproduction. (**A-C**) IIS activity is represented by phosphorylation of AKT and MAPK (P-AKT and P-MAPK), normalized by total AKT and MAPK (T-AKT and T-MAPK). (**A**) Western blot analysis of P-AKT, T-AKT, P-MAPK and T-MAPK in the fat body (left) and different parts of the dissected ovary (right: germarium, early egg chamber (EC) and late EC tissues from worker (W) *vs*. gamergate (G). (**B**) Quantification of the relative P-AKT and P-MAPK levels in the fat body of W and G described in (A). p values are from unpaired t test (p<0.01**, p<0.001***, n=6). Bars and error bars are represented mean ± SEM. (**C**) Immunofluorescence (IF) staining of transcription factor FOXO in the fat body as detected by *H. saltator* FOXO antibody (green) and DNA staining by DAPI (blue). Left panel: FOXO localization in the cytoplasm of worker fat body. Right panel: FOXO localization in the nucleus of gamergate fat body. Nuclei were identified by DAPI and indicated by arrows in the FOXO staining. (**D**) Averages of the yolky oocyte number per workers and RT-qPCRs for *vitellogenin* (*vg*) mRNA in abdominal fat body of workers 7 days post-injection with either DMSO or U0126 at different dosages (10, 100 and 1,000 μM). Representative bright-field images of dissected ovaries (middle panels). n=3 independent colonies. Bars and error bars represent mean ± SEM. p-values are from Kruskal-Wallis test with multiple comparisons. *Rpl32* is used as a reference gene for data normalization. Data are from more than 10 biological replicates per condition.

Likewise, using phosphorylated AKT as a readout (Fig. S2D), we analyzed IIS-AKT activity in multiple tissues in gamergates *vs*. workers. Strikingly, the gamergate fat body had a significantly lower activity of IIS-AKT compared to workers (approximately 2x decrease, n=6, p<0.01, Fig. 3A, B). This is consistent with evidence that reducing insulin signaling in the fat body (which includes adipose tissue) lengthens the lifespan of other species, *e*.*g*. worms, flies and mice (*14*). In contrast, IIS-AKT activity was upregulated in the PPG of gamergates, and was activated at similar levels in the brain and the MV of workers and gamergates (Fig. S5E). as well as in the germarium (Gm), the ovarian region containing the germline stem cells in workers and gamergates (Fig. 3A). However, AKT phosphorylation was very low in the ovary in early- and late-stage egg chambers that are only present in gamergates (Fig. 3A). We also analyzed the subcellular localization of FOXO in these tissues. An inactive IIS pathway results in unphosphorylated FOXO being localized to the nucleus, but activation of the IIS pathway results in FOXO phosphorylation by AKT, leading to its retention in the cytoplasm (*23*). In accordance, FOXO was localized in the nucleus in the gamergate fat body, whereas it was localized in the cytoplasm of workers (Fig. 3C). In the gamergate ovary, FOXO was localized in the nucleus in all follicle cells and in the nurse cells of egg chambers up to stage 4, while at stage 5, FOXO was only localized to the cytoplasm in nurse cells (Fig. S5F), suggesting higher IIS-AKT activity in late-stage nurse cells. One of the target genes that is transcriptionally repressed by FOXO, *4-hydroxyphenylpyruvate dioxygenase* (*Hpd-1)*, encodes an enzyme involved in the degradation of tyrosine, the amino acid precursor for biogenic amines (*56*). *Hpd-1* was significantly downregulated in the fat body of gamergates compared to workers (Fig. 2B). In *C. elegans* and *Drosophila*, whole-body knockdown of *Hpd-1* leads to extended lifespan, which was attributed to increased levels of dopamine and octopamine that have a protective role in neuromuscular functions (*56, 57*). Interestingly, the levels of dopamine but not those of octopamine were significantly higher in *Harpegnathos* gamergates compared to workers and their levels positively correlated with ovary development (*58*). Thus, *Hpd-1* downregulation, which correlates with reduced IIS-AKT activity in the gamergate fat body, might additionally explain the extended longevity of reproductive gamergates.

Taken together, we conclude that Ins levels are increased in the gamergate brain, leading to the activation of the MAPK pathway in the fat body and ovary. However, AKT activity is reduced in the fat body and part of the ovary of gamergates. These results raise two questions: does MAPK, but not AKT, lead to the production of vitellogenin to promote ovary growth and egg production, while decreased AKT activity in the gamergate fat body presumably leads to extended longevity? If so, how is AKT (but not MAPK) downregulated in the gamergate?

### MAPK activity is required for ovary growth

We (Fig. 2E, F), and others (*47*) have shown that Ins induces reproduction in ants. Since MAPK, but not IIS-AKT, is active in the gamergate fat body and ovary, we tested the role of IIS-MAPK on caste transition and ovary activation using the U0126 MEK inhibitor, which prevents MAPK phosphorylation. Without a supplemented Ins peptide, the inhibitor was able to specifically inhibit MAPK phosphorylation in the dissected fat bodies of workers (3X decrease in treated fat body compared to control), but it did not affect AKT phosphorylation (80% of activity remained in treated tissue compared to control). Upon co-treatment with the Ins peptide, MAPK phosphorylation was reduced by 2X with U0126, while AKT phosphorylation remained almost unchanged (90% of control activity, Fig. S5G).

We also injected U0126 in workers during their transition to gamergates and measured *Vitellogenin* expression as a molecular marker for egg production and scored ovary growth 6 days after injection. Inhibition of MAPK phosphorylation by U0126 led to a concentration-dependent decreased expression of *Vitellogenin* in the fat body as well as the number of yolky oocytes in dueling workers (Fig. 3D). While oogenesis was promoted by Ins injection to workers (Fig. 2E, F), the co-injection of U0126 with Ins impeded this effect (Fig. 2G). Interestingly, the germarium remained intact, suggesting that the inhibitor did not cause atrophy of the germline stem cell niche, which is present in both workers and gamergates.

Based on these results, we conclude that Ins from the brain can activate the MAPK pathway in the fat body and ovary and, as a result, induce the production of mature egg chambers. MAPK activation is also necessary to promote *vitellogenin* expression in the fat body, as well as ovary growth and the transition to being reproductive. In contrast, IIS-AKT activity is decreased in the fat body and the developed ovaries of gamergates, but remains comparable in the germarium of workers and gamergates. The constant activity of IIS-AKT in the germarium of both castes likely plays an important role in germline stem cell maintenance and early differentiation, as it does in *Drosophila* and *C. elegans (59-61)*. In this manner, workers can retain a functioning germline that will allow them to reactivate oogenesis if they ever become gamergates.

### The IIS inhibitors, Imp-L2 and ALS, are upregulated in the ovary

To explain the paradox of upregulation of Ins and IIS-MAPK and downregulation of IIS-AKT activity in gamergates, we searched for candidate genes that could mediate the decreased IIS-AKT activity in the fat body and some ovarian tissues. In our differential expression analysis, *insulin receptor 1 and 2* (*InR1/2*) were both down-regulated in the gamergate fat body and whole ovary (Fig. 2B). Additionally, two genes encoding secretory proteins that inhibit the IIS pathway were upregulated in the ovary of gamergates: *Imaginal morphogenesis protein-Late 2* (*Imp-L2*) and *Acid-labile Subunit* (*ALS*) (*62-64*) (Fig. 4A). In *Drosophila*, Imp-L2 and ALS, either individually or together, bind to circulating Dilp2 and Dilp5, the Ins homologs produced in brain IPCs that are secreted into the hemolymph. Imp-L2 and ALS antagonize IIS in peripheral tissues (*16, 39, 62-65*). Likewise, in mammals, ALS and IGF-binding protein 7 (IGFBP7), a homolog of Imp-L2, both bind plasma insulin and IGF and subsequently restrain their interaction with their receptors (*66-69*). IGFBP7 displays higher binding affinity to insulin than to IGF (*67*). This evidence suggests that the elevated expression of *Imp-L2* and *ALS* genes in gamergate ovaries and their secretion into the hemolymph antagonize the IIS-AKT pathway in both mature ovarioles and the fat body.

**Fig. 4.**
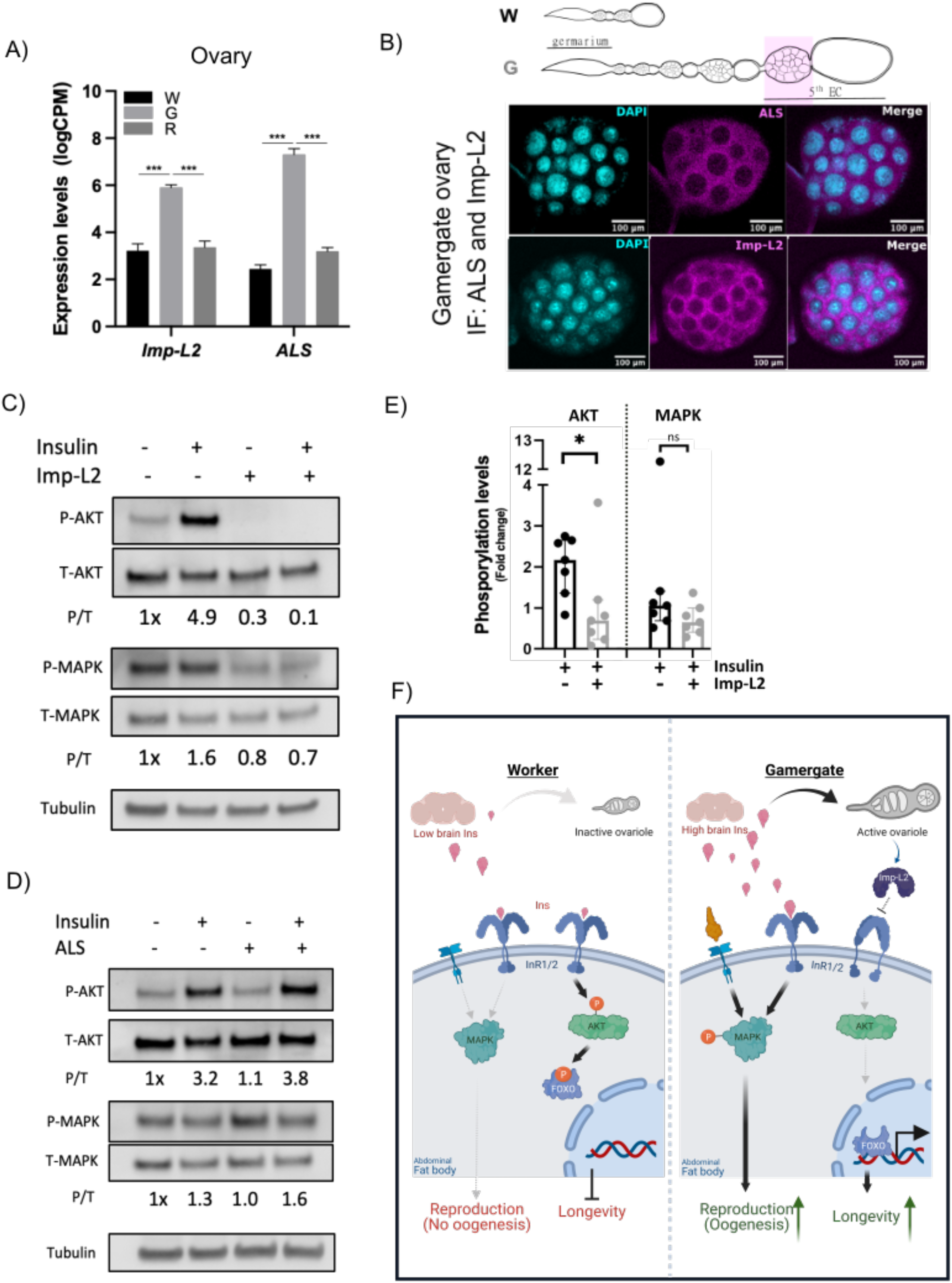
Imp-L2 produced from gamergate ovaries preferentially antagonizes the IIS-AKT pathway. (**A**) RNA-seq levels of ovarian *ALS* and *Imp-L2* in workers (W), gamergates (G) and revertants (R). Bars and error bars represent mean ± SEM. (**B**) IF staining of insulin/IGF-binding proteins (ALS and Imp-L2) in the late stages of the egg chambers of a gamergate ovary as indicated by a pink box in the schematic on top. Magenta represents ALS and Imp-L2 in upper and lower panels, respectively. DAPI is shown in cyan. (**C-D**) Western blot analyses showing the effects of recombinant Imp-L2 or ALS (produced in SF9 cells) on IIS-AKT and -MAPK pathways in the abdominal fat body tissue dissected from the candidate worker. Levels of P-AKT, T-AKT, P-MAPK, T-MAPK and tubulin detected in the fat body as a function of treatment with insulin peptide (Ins) minus/plus recombinant Imp-L2 (**C**) or ALS protein (**D**). (**E**) A quantitative plot of phosphorylation levels of AKT and MAPK from worker fat bodies incubated with or without Imp-L2 in the presence of insulin peptide. Bars and error bars represent median with interquartile range. p-values are from Mann-Whitney test (p<0.05*, n=7 biological replicates). (**F**) Proposed models of reproduction-associated longevity in ants. Left: a shorted-lived worker; right: a long-lived gamergate. The basal level of *insulin* (*Ins*) secreted from the worker brain is sufficient to activate the IIS-AKT pathway via insulin receptors (InRs) in the abdominal fat body, resulting in FOXO phosphorylation and a normal lifespan in workers. In contrast, the high level of *Ins* in the gamergate brain promotes ovary maturation and contributes to MAPK activation in the fat body. An unidentified receptor (left to InR1/2) may also contribute to MAPK activation. Additionally, the gamergate ovary produces Imp-L2 protein that antagonizes IIS-AKT signaling in the fat body, giving rise to nuclear localization of FOXO and longevity in gamergates.

Major sources of ALS in mammals are the liver (*70*), kidney and ovary (*71, 72*), while *Imp-L2* mRNA is found in ovarian follicle cells (*73*). To further test the expression profiles of *Imp-L2* and *ALS* genes in *Harpegnathos*, we performed RT-qPCR on multiple tissues, including the germarium in the ovary, early and late-stage egg chambers and yolky oocytes. Indeed, *Imp-L2* and *ALS* mRNA were mainly expressed in ovaries, especially in late-stage egg chambers and yolky oocytes that are only present in gamergates. In contrast, their expression levels were relatively low in other tissues, such as the brain, fat body, gut and early-stage egg chambers (Fig. S6A). We generated antibodies against Imp-L2 and ALS proteins. These two proteins were only detected in late-stage egg chambers (Fig. 4B and S6B, C). These results strongly suggested that both IIS inhibitors, Imp-L2 and ALS, are predominantly expressed in gamergate ovaries, in particular, in well-developed egg chambers from which they are secreted to act on abdominal tissues, including the fat body.

### Imp-L2 specifically blocks AKT in the fat body

*Imp-L2* and *ALS* expressed in the ovary may act as inhibitors that contribute to decreased IIS-AKT activity in the fat body and ovary of gamergates, thereby increasing lifespan. To test whether these proteins affect IIS in ants, we generated FLAG-tagged versions of *Harpegnathos Imp-L2* and *ALS* in baculovirus and purified the proteins from lysates of transfected SF9 insect cells using FLAG-based affinity purification (Fig. S6D). These purified proteins were then used for *ex vivo* assays using dissected abdominal fat body tissues from workers. To minimize individual variations, dissected fat body tissue from the same individual was separated for co-incubation with Ins, with or without Imp-L2 or ALS. Imp-L2 was able to strongly inhibit activation of IIS-AKT (3x reduction compared to control, p<0.05, n=7, Fig. 4, C-E), but ALS was ineffectual (Fig. 4D). In contrast, IIS-MAPK was not significantly inhibited by either Imp-L2 (Fig. 4, C-E) or ALS (Fig. 4D), suggesting that Imp-L2 exhibits a specific role in downregulating IIS-AKT, but not IIS-MAPK in gamergates. In support of our results, a study of human IGFBP7 shows that IGFBP7, a homolog of Imp-L2, negatively regulates IIS-AKT signaling, but has no effect on the IIS-MAPK pathway (see figure 1C in ref (*74*)).

Thus, our findings support the model that Imp-L2 produced by the ovary of gamergates acts as an Ins inhibitor to specifically reduce IIS-AKT activity in the fat body and ovary, thus extending lifespan. In contrast, IIS-MAPK is unaffected and appears to be the primary regulator for initiating and sustaining the reproductive function of the ovary, in particular by producing vitellogenin and perhaps mobilizing lipids that will accumulate in the egg. The modulation of IIS-AKT vs. IIS-MAPK through activation and inhibition offers an effective solution to the discrepancy between increased insulin and reproduction and prolonged lifespan (Fig. 4F).

## Discussion

The traits that favor reproduction have particular importance in eusocial insects as the reproductive duty of the whole colony is placed on one or very few queens that are highly prolific. At the same time, such individuals are difficult or impossible to replace without disrupting the colony, so they must have a very long lifespan to allow the colony to thrive beyond the individual life of its workers (*28, 75*). Natural selection has thus co-opted molecular pathways that increase lifespan in queens. Here, we described how in *Harpegnathos*, systemic activation of the IIS pathway driven by the production of Ins in the brain facilitates active reproduction via IIS-MAPK, but is combined with inhibition of IIS-AKT signaling in the fat body and developed ovary, thereby extending lifespan. This local effect correlates with lower expression of InRs in the fat body and ovary and appears to result from the inhibitory effect of Imp-L2 secreted from the reproductive ovary.

The highly conserved IIS-AKT pathway is responsible for sugar uptake and lipid metabolism in insects and vertebrates (*76-78*): it promotes glycogen synthesis and suppresses lipogenesis via phosphorylation of the downstream metabolic factor FOXO (*79, 80*). For example, reduced IIS-AKT and nuclear FOXO promote *fatty acid synthase (Fasn)* expression in the mouse liver (*81, 82*). In accordance, the upregulated expression of *Fasn* that we observed in the gamergate fat body must be due to reduced IIS-AKT activity. In addition, reduced glucose and trehalose in the fat body of gamergates might lead to increased lipogenesis as carbohydrate breakdown through glucose-pyruvate conversion provides acetyl-CoA precursors for fatty acid synthesis (*83, 84*). However, we observed decreased triglyceride storage in the fat body and increased circulating triglyceride in the hemolymph. We speculate that newly synthesized lipids are not stored in the fat body but are redirected through the hemolymph to the ovary to promote egg production in gamergates. As gamergates feed constantly, that energy storage via lipid is less important. Instead, lipids are mainly utilized by the ovary for continuous oogenesis. In parallel, the volume of the brain that is an organ with high energy demand, is reduced by ∼20% in gamergates compared to workers (*36, 85*), thereby saving energy to maximize reproduction.

As in raider ants (*47*), the regulation of reproduction in *Harpegnathos* gamergates is achieved by increased insulin production. However, the peculiar lifestyle of raider ants (*47*) does not require an extended lifespan. In order to foster the extended lifespan of gamergates, the increased insulin in *Harpegnathos* does not stimulates the IIS-AKT pathway, but instead, only the IIS-MAPK branch. Indeed: (1) Ins production is elevated in the brain, (2) IIS-MAPK activity is elevated in the fat body and ovary, (3) Ins treatment promotes vitellogenesis and oogenesis, and (4) inhibition of MAPK activity reverses these effects. Our findings suggest that IIS-MAPK is required for the maturation of egg chambers and consequently the development of oocytes. In accordance, it has been argued that in *Drosophila*, vitellogenesis is not regulated by IIS-AKT/FOXO, but via the IIS-MAPK branch (*49-51, 86*). In mice, the exercise hormone Irison induces MAPK signaling, which mediates a conversion of white (energy-storing) adipocytes into brown (energy-consuming) adipocytes, thereby reducing lipid storage (*87*). This phenomenon may explain the correlation between MAPK activation and increased vitellogenesis or lipid utilization in the gamergate fat body as was observed by the changes in fat body structure and decreased lipid content. Additionally, our data demonstrates that Ins mildly activates MAPK activation, while MAPK activity in the gamergate fat body is ∼4x higher than in workers; other ligands, in addition to Ins, may be responsible for MAPK activation in gamergates. Interestingly, we had shown that gamergates exhibit increased ecdysone titers (*35, 88*) as well as MAPK activation (Fig. 3A, B), which might form a positive feedback loop. This scenario is reminiscent of *Drosophila* wherein ecdysone promotes intestinal growth via increased EGFR ligands leading to activation of the MAPK pathway (*89, 90*), which is itself required for ecdysone production (*91*). Taken together, we suggest a model in which the worker transition to gamergate is accompanied by Ins activation of IIS-MAPK, which triggers vitellogenin expression in the fat body and reactivates ovary growth. The follicle cells in the growing egg chambers produce ecdysone, further enhancing MAPK activity, oogenesis and reproduction. In *Drosophila*, Imp-L2 was identified as an ecdysone-inducible gene (*92*); therefore, ecdysone might also trigger the expression of anti-insulin Imp-L2 in the gamergate ovary.

We found that IIS-AKT activity is reduced in the fat body and early-stage egg chambers (stages 1-4). This appears to be due to the production of Imp-L2 from the mature ovary, which is secreted into the hemolymph and inhibit Ins-induced AKT activation in the fat body. However, some tissues seem resistant to the effect of Imp-L2. For example, the germarium shows similar AKT activity in both workers and gamergates, consistent with the role of IIS-AKT in germline stem cell maintenance and early differentiation in *Drosophila*. In addition, the post pharyngeal gland (PPG) in the gamergate head shows an increase in phosphorylated AKT. As the PPG is involved in the storage of lipids and cuticular hydrocarbons (CHCs) (*54, 55*), higher IIS-AKT activity may be essential for the production of gamergate-specific CHC pheromones to facilitate social recognition (*93*). In the ant *Myrmecia gulosa*, CHC extracts from the PPG of reproductive females are more attractive to their nestmates than those from non-reproductive workers, while extracts from other pheromone-producing glands, such as Dufour and mandibular glands, do not show this effect, indicating a specific role of the PPG content as queen pheromones (*94*).

The question therefore arises as to how do some tissues protect themselves from anti-insulin inhibition by Imp-L2? One protective candidate is the membrane-bound serine protease matriptase found in mammals. Matriptase proteolytically cleaves IGFBP7/Imp-L2 protein, thereby preventing local IIS-AKT suppression, while showing no effect on IIS-MAPK in mammals (*74, 95*). In *Drosophila*, Notopleural is reported to be a functional homolog of human matriptase (*96*), but its function in regulating IIS activity is not clear. We speculate that matriptase/Notopleural provides a protective role against the Imp-L2 effect in the tissues that produce it, *e*.*g*. the germarium, thereby maintaining normal IIS function in gamergates. Alternatively, other pathways might also provide such a protective role. For example, intestinal tumors in *Drosophila* produce Imp-L2 as a wasting factor that cause IIS-AKT reduction and atrophy of surrounding tissues (*97-99*). However, tumor cells are resistant to Imp-L2 due to increased Wingless (Wg) that opposes the Imp-L2 effect and maintains tumor growth (*100*). A similar self-evasion mechanism via Wg or other pathways might also exist in other gamergate tissues, such as the ovary and PPG.

In summary, our study reveals a mechanism to retard aging and achieve longevity in ants. Based on the hyper-function theory of aging, the activity of processes required for reproduction in early stages of life shortens lifespan as they retain a sustained activity in later stages (*101, 102*). Ants provide a unique solution to restrict hyperactivity throughout their very long reproductive life via selective inhibition of IIS-AKT by Imp-L2. Through crosstalk of IIS components in multiple tissues, IIS-MAPK and IIS-AKT differentially regulate ovary growth, germline maintenance and lifespan. This interplay, which evolved in ants and perhaps in other eusocial insects, reconciles the increased longevity associated with reproduction.

## Acknowledgements

We thank New York Langone Health (NYULH) Genome Technology Center for help with sequencing; Y. Deng and the NYULH Microscopy Core for help with confocal imaging; H. Yang, J. Gospocic, C. Penick, K. Haight and J. Liebig for their insightful suggestions and technical support; and the entire Desplan and Reinberg laboratories for helpful discussions; and L. Vales for help with manuscript preparation.

## Funding

This work was supported by HHMI Collaborative Innovation Award (CIA) #2009005 and HCIA #2009005 to D.R. and C.D.; NIH grants R21GM114457 to D.R.; R01EY13010 to C.D.; R01AG058762 to D.R. and C.D.; NIH Ruth L. Kirschstein NRSA postdoctoral fellowship F32AG044971 and NSF I/UCRC CAMTech grant IIP1821914 to H.Y.; Human Frontier Science Program long-term fellowship LT000010/2020-L to B.S.

## Author contributions

H.Y., C.O., F.C.-A., G.M., C.D., and D.R. designed the study; H.Y., C.O., and G.M. performed transition experiments, lifespan monitoring and sample collection; C.O., and N.D. performed bioinformatic analyses; H.Y., C.O., G.M., A.L., and B.S. performed ISH/IF staining; C.O., and J.M. injected ants; H.Y., C.O., F.C.-A., J.M., and L.D. performed insulin and pharmacological experiments; X.Z. performed phylogenetic analysis; H.Y., C.O., F.C.-A., J.M., and M.T. quantified IIS activities; F.C.-A. performed ALS and Imp-L2 assays; H.Y., C.O., and F.C.-A. wrote the manuscript with feedback from all authors; C.D., and D.R. supervised the project.

## Competing interests

The authors declare no competing interests.

## Data and materials availability

Raw sequence data are available through NCBI (BioProject: TBD).

## Supplementary materials

**Fig. S1.**
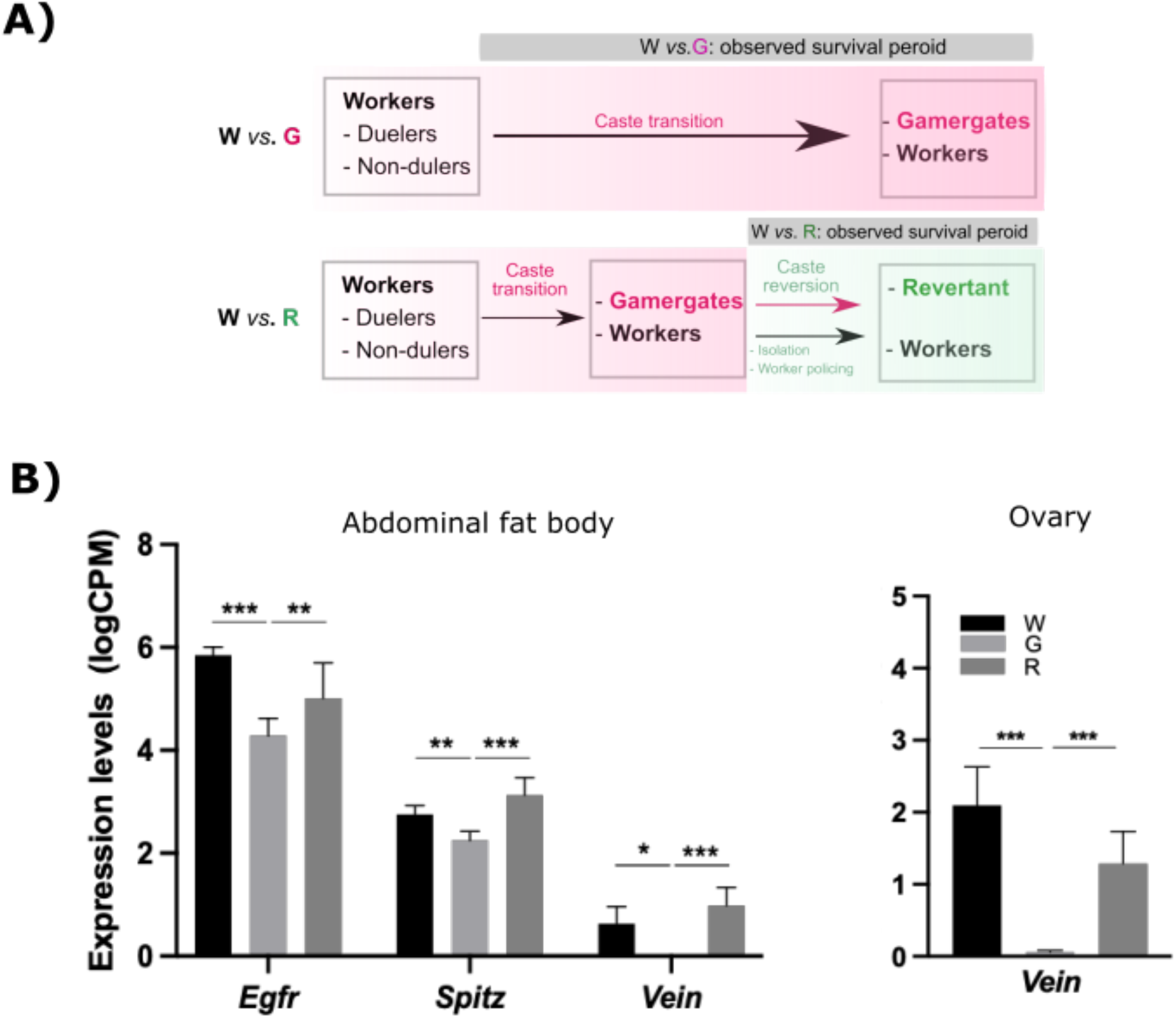
Diagram of the timeline for caste transition [Worker (W) to Gamergate (G)] and reversion [Gamergate (G) to non-reproductive revertant (R)] and differentially expressed genes (DEGs) between W *vs*. G in the brain, abdominal fat body and ovary. (**A**) A colony consisting of 30 similarly aged matched workers initiates the dueling tournament, in which workers can be classified into two groups (dueling and non-dueling). The caste fate is determined based on dueling activity and oviposition. Survival rates of worker (W) and gamergate (G) are observed during the transition for over a year. For the process of G-to-R reversion, mature gamergates derived from an approximate 3-month transition, along with their non-reproductive nestmates are subjected to reversion by individual isolation followed by worker policing in an established queen or gamergate colony. The survival data of revertants and non-reproductive nestmates are collected during the reversion process. Individuals used in the reversion (lower image) are independent from the W-to-G transition (upper image). (**B**) RNA-seq levels of DEGs in the abdominal fat body and ovary in workers (W, black), gamergates (G, grey) and revertants (R, dark grey). *Egfr: epidermal growth factor receptor*. Data are from four biological replicates per caste. p values from EdgeR are indicated. Bars and error bars represent mean ± SEM. False Discovery Rate (FDR) cut off < 0.05.

**Fig. S2.**
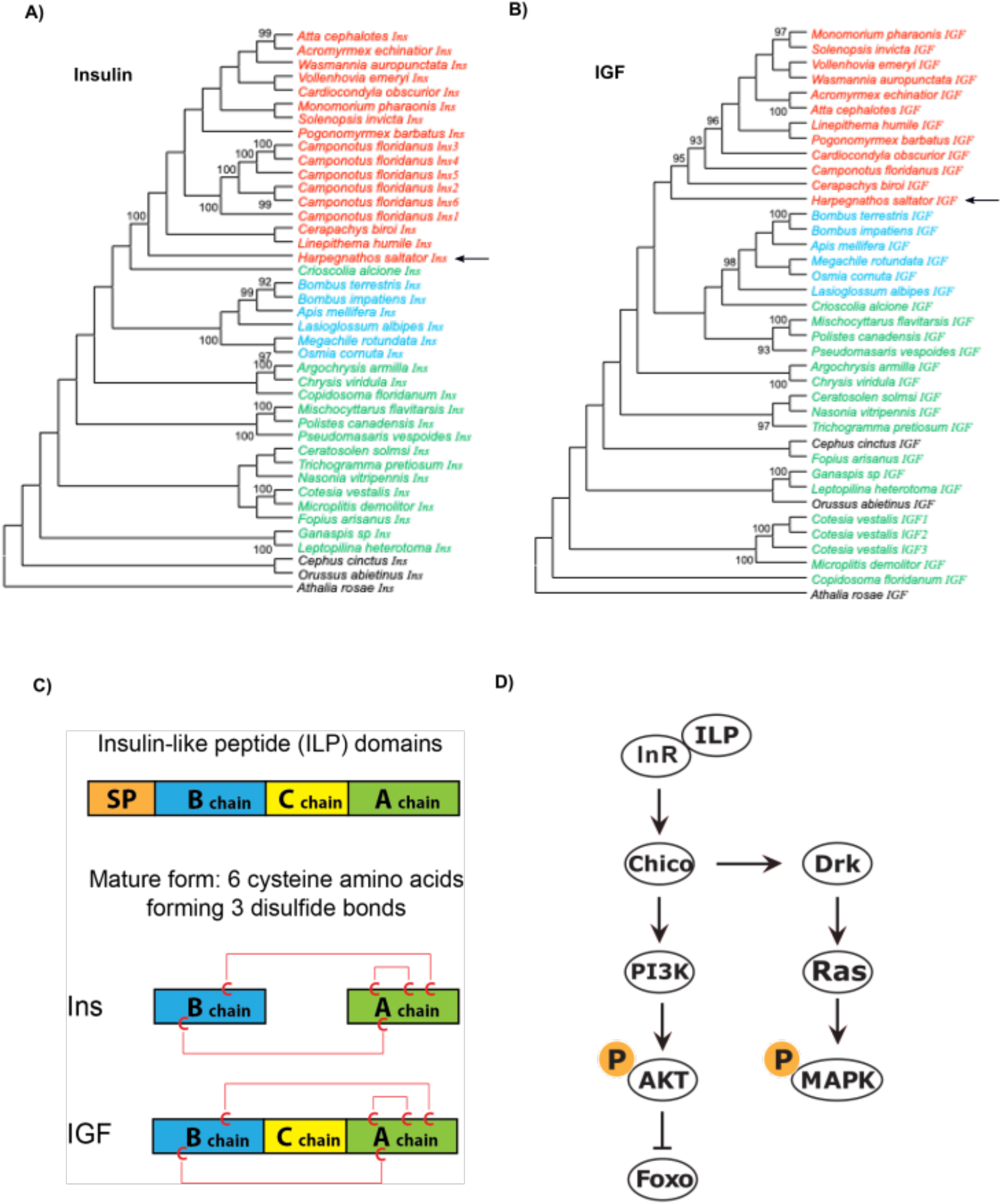
Phylogenetic trees and structures of Ins and IGF. (**A-B**). Phylogenetic trees of *hymenopteran Ins* (**A**) and *IGF* (**B**) genes. Genes from ants, bees, wasps, and sawflies are indicated in red, blue, green, and black font respectively. Numbers on internal branches represent support values as measured by the Ultrafast Bootstrap approach with 1000 replicates. Support values lower than 90% are not shown. (**C**) Domain structures in insulin-like peptides (ILPs) and the mature forms of Ins and IGF proteins. Red lines show disulfide bonds. SP: signal peptide. C: cysteine. (**D**) Highly conserved Insulin/IGF signaling pathway (IIS): the insulin/IGF signaling (IIS) pathway and its downstream targets, AKT and MAPK. AKT and MAPK can also be activated by IIS-independent factors. P: phosphorylation.

**Fig. S3.**
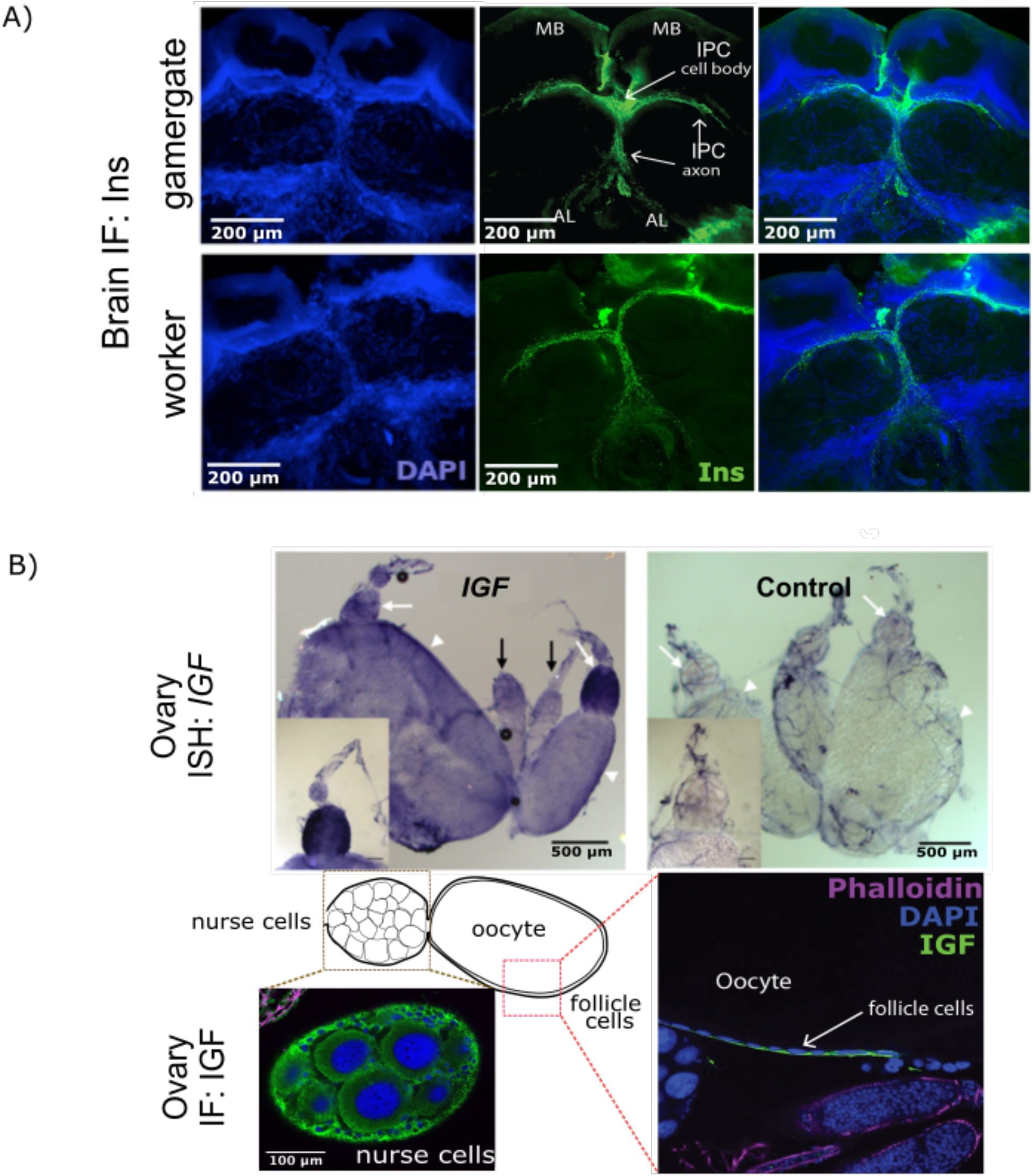
Localizations of *IGF and Ins* in ovary and the brain. (**A**) immunofluorescence (IF) staining of brain insulin-like peptide (Ins) in a gamergate (upper panel) and a worker (lower panel). Ins peptide (green) localizes near insulin producing cells (IPCs) situated between two mushroom bodies (MBs), and along the axons in both brains of worker and gamergate. Blue represents DAPI. (**B**) Colorimetric *in situ* hybridization (*ISH*) and IF staining of ovarian IGF. Upper panel: Localization of *IGF* mRNA by *ISH* as indicated by arrows in top-left panel. Top-right panel shows a control probe. Lower panel: schematic displaying the nurse cell and the developing oocyte in an egg chamber (EC, left panel) and localization of IGF protein in the EC (right and lower panels). Ovarian IGF protein (green) localizes in the cytoplasm of follicle cells (right panel) and nurse cells (lower-left panel). DAPI stains for DNA (blue). Phalloidin is represented by magenta.

**Fig. S4.**
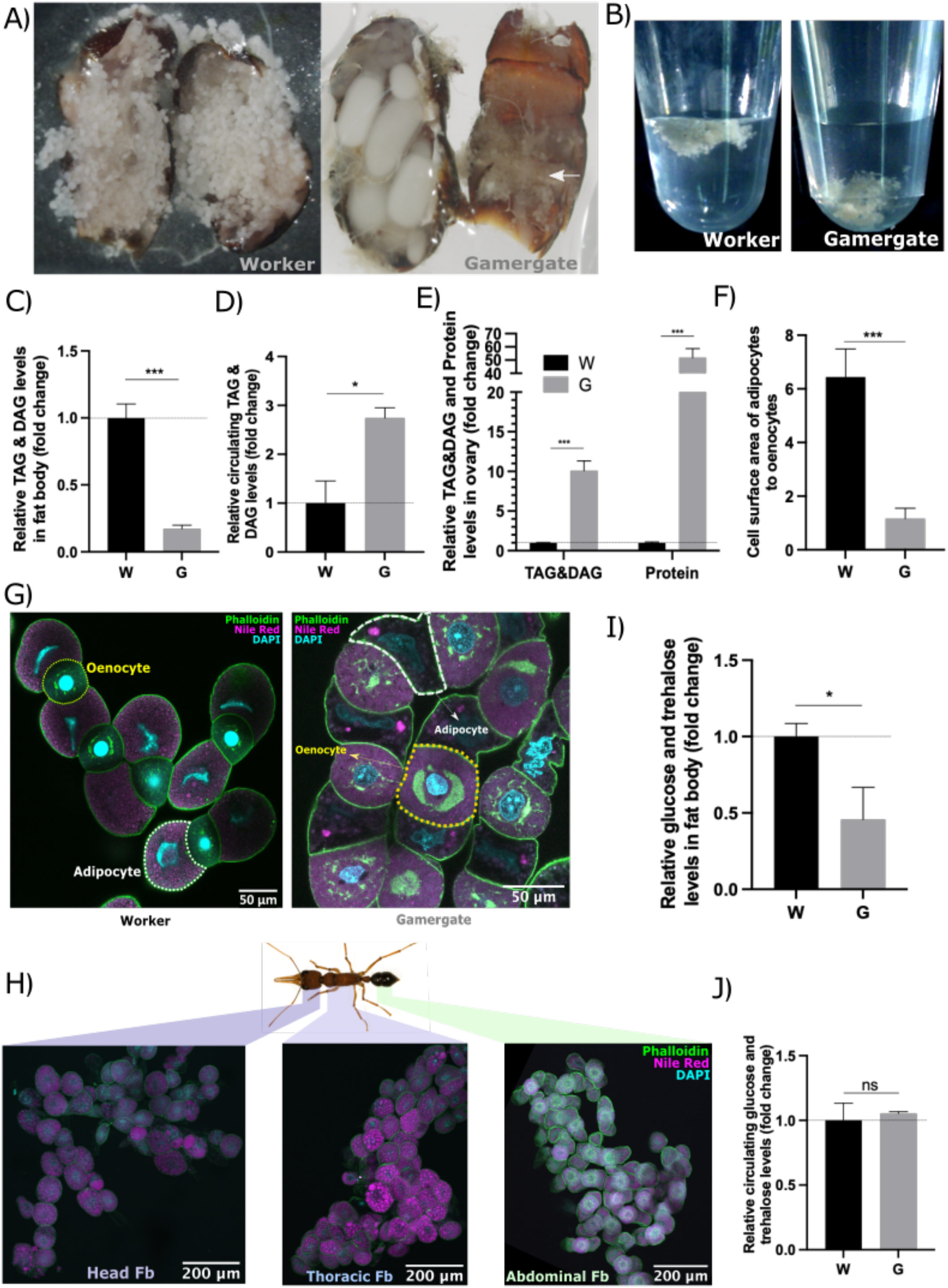
Lipid depletion in the abdominal fat body during the worker-to-gamergate transition. (**A-B**) Morphological comparisons of abdominal fat bodies in worker vs. gamergate. (**A)** Dissected abdomens of workers (W, left) and gamergates (G, right) were imaged by a bright field microscope. Arrow in G indicates fat body. (**B**) The dissected abdominal fat bodies from workers (left) and gamergates (right) float and sink in saline buffer, respectively. (**C-E**) Comparison of the triglyceride (TAG) and diglyceride (DAG) levels in the fat body (C), hemolymph (D) and ovary (E) of worker and gamergate. (**C**) Quantification of total amounts of TAG and DAG in abdominal fat body per individual worker (W) vs. gamergate(G) (n=5, p<0.001***). (**D**) Quantification of circulating TAG and DAG levels per volume (μl) in worker vs. gamergate hemolymph (n=3, p<0.05*). (**E)** Quantification of TAG and DAG and protein levels in whole ovaries from individual workers (n=10) and gamergates (n=8, p<0.001***). (**F-H**) Immunofluorescence (IF) staining of the fat body with DAPI, Nile Red and fluorophore-conjugated Phalloidin antibody (cyan, magenta and green, respectively). (**F**) Quantification of the ratio of adipocyte to oenocyte cell surface area in workers vs. gamergates (n=3 biological replicates, p<0.001***). p values are from unpaired t test. Bars and error bars represent mean ± SEM. (**G**) Two fat body cell types: adipocytes (white borderline) and oenocytes (yellow borderlines), are indicated in the abdominal fat body of both worker (left) and gamergate (right). (**H**) Oenocytes or hepatocyte-like cells, recognized by a green ring structure detected by Phalloidin antibody, are found exclusively in the abdominal fat body (Fb), and not in the head and thoracic fat bodies. (**I-J**) Comparison of the sugar levels (glucose and trehalose) in worker vs. gamergate. (**I**) Quantification of total sugar levels in the abdominal fat body of each worker vs. gamergate (n=5, p<0.05*). (**J**) Quantification of circulating sugar levels (per microliter) in worker vs. gamergate hemolymph (n=3, p=0.70). p values are from unpaired t test. Bars and error bars represent mean ± SEM.

**Fig. S5.**
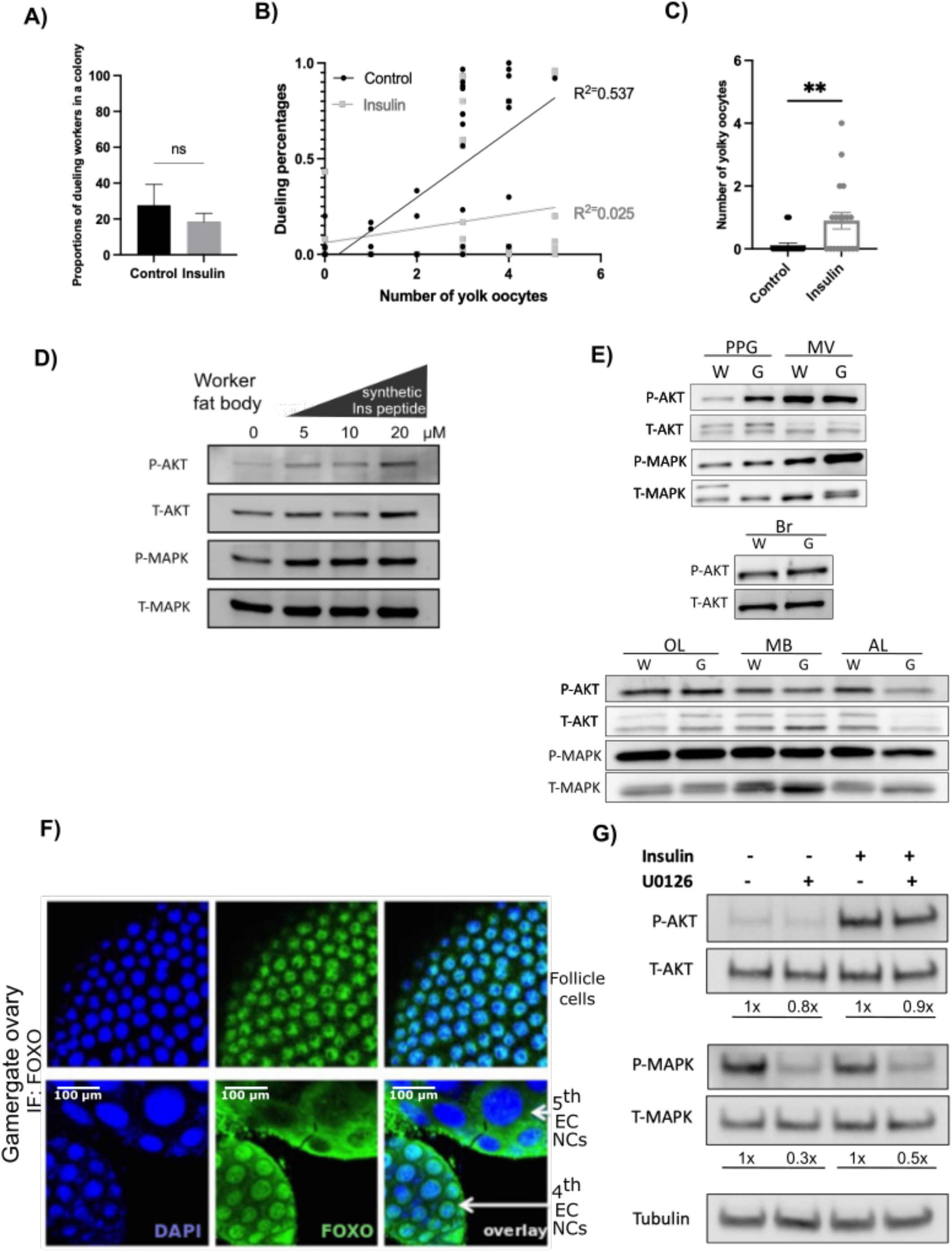
Effects of insulin and U0126 on antennal dueling in queenless workers. (**A-B**) A single injection of insulin and its solvent into worker abdomen. (**A**) The proportion of dueling vs. non-dueling workers in a colony are not significantly affected by Ins injection (**B**) Positive correlation between dueling activity and ovary development (the number of yolky oocytes) in control workers (black, R^2^=0.5), as opposed to insulin-injected workers (grey, R^2^=0.02) that exhibit no positive correlation. (**C**) The number of yolky oocytes of workers in a mature colony with reproductives, injected with mock or insulin, were estimated 5 days post injection (n=3 colonies, 60 individuals per condition, p<0.01**). p values are from Mann-Whitney test. Bars and error bars represent mean ± SEM. (**D**) Western blot analysis of phosphorylated AKT (P-AKT), total AKT (T-AKT), phosphorylated MAPK (P-MAPK) and total MAPK (T-MAPK) levels in worker fat bodies treated with different concentrations of synthetic Ins peptide (0, 5, 10 and 20 μM). (**E**) Western blot analysis comparing P-AKT, T-AKT, P-MAPK and T-MAPK levels in the following tissues from worker (W) vs. gamergate (G): the postpharyngeal gland (PPG), malpighian vesicle (MV), the whole brain (Br), optic lobe (OL), mushroom body (MB) and antennal lobe (AL). (**F**) IF staining of a transcription factor FOXO in the gamergate ovary detected by *H. saltator* FOXO antibody. Nuclear-localized FOXO in the ovarian follicle cells and in the 5^th^ egg chamber (EC) nurse cells (NCs) are shown in upper and lower panels, respectively. FOXO localized in the cytoplasm is shown in the 4^th^ EC NCs (lower panels). FOXO represents in green. Nuclei were identified by DAPI (blue). (**G**) Western blot analysis of P-AKT, T-AKT, P-MAPK, T-MAPK and tubulin in worker fat body treated with the synthetic Ins peptide and/or U0126. Fold changes (X) of U0126-treated samples are indicated in comparison to either control or Ins-treated samples.

**Fig. S6.**
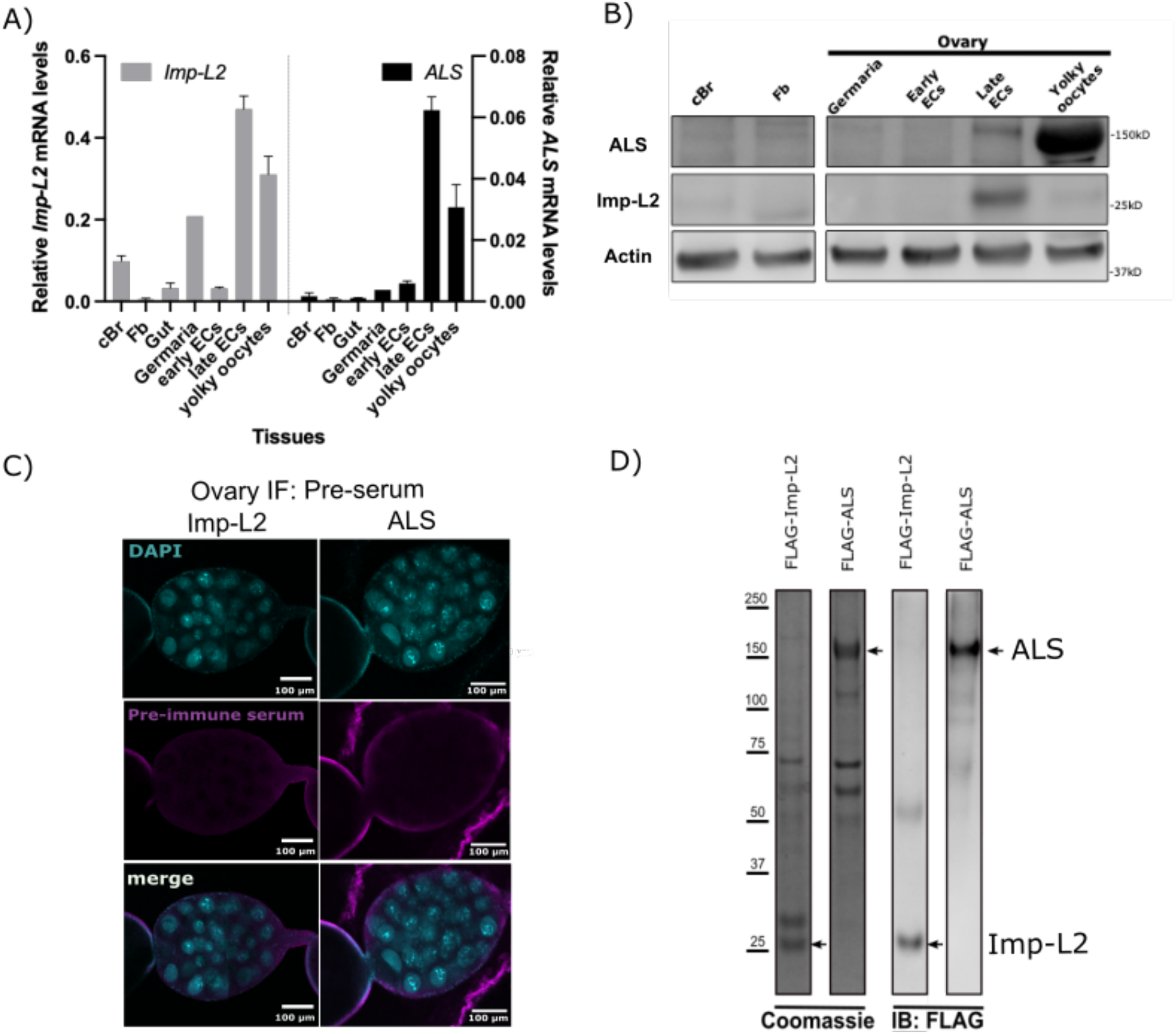
Localization of FOXO, Imp-L2 and ALS in gamergate ovaries. (**A**) RT-qPCR analysis of *ALS* and *Imp-L2* mRNAs in different tissues, such as the central brain without optic lobes (cBr), fat body (Fb), gut and different parts of the ovary from gamergates, including germaria, early egg chambers (ECs), late ECs and yolky oocytes (n=3). Germaria from three individuals were pooled together. Expression levels were normalized by *ribosomal protein L32 (Rpl32)* gene. Bars and error bars represent mean ± SEM. (**B**) Western blot analysis of ALS and Imp-L2 proteins from different tissues, as indicated. Actin serves as a loading control. (**C**) IF staining of *H. saltator* ALS and Imp-L2 pre-sera in the gamergate ovary (magenta). (**D**) Purified FLAG-tagged Imp-L2 and ALS proteins from insect Sf9 cells were separated by SDS-PAGE and stained with Coomassie Blue (left) and immunoblotted with FLAG antibody (right).

**Table S1.** Information for gene identifiers of *Harpegnathos saltator* genes discussed in the text.

| Used in this study |  | Hsal gene ID version 8.5 | NCBI Reference Sequence<br>(Accession number.Version) | Tissue sources | Gene ID |
| --- | --- | --- | --- | --- | --- |
| Gene symbol | Gene description |  |  |  |  |
| *Ins* | Insulin or LIRP | HSALG004952.1 | XM_011149533.3 | Brain | LOC105188195 |
| *Crz* | corazonin | HSALG005445.2 | XM_011155750.2 | Brain | LOC105191954 |
| *InR1* | Insulin receptor 1 | HSALG011112.2 | XM_025302516.1 | Abdominal fat body | LOC105185958 |
| *InR2* | Insulin receptor 2 | HSALG009512.2 | XM_019841795.2 | Abdominal fat body, Ovary | LOC105183944 |
| *Vg* | vitellogenin | HSALG009096.2 | XM_011138688.2 | Abdominal fat body | LOC105181726 |
| *IGF* | insulin-like growth factor I | HSALG003795.2 | XM_011147513.3 | Ovary | LOC105186969 |
| *ALS* | Acid-labile subunit, convoluted, IGF binding protein (IGFBP), protein artichoke | HSALG003656.2 | XM_011151816.3, NM_001299487.1 | Ovary | LOC105189593 |
| *Imp-L2* | Imaginal morphogenesis protein-Late 2, Ecdysone-inducible gene L2 | HSALG000642.2 | XM_011141551.3, NM_079197.5 | Ovary | LOC105183446 |
| *Hpd-1* | 4-hydroxyphenylpyruvate dioxygenase | HSALG007876.2 | XM_011141398.3, NM_168857.3 | Abdominal fat body | LOC105183352 |
| *Far* | fatty acyl-CoA reductase CG5065, CG5065 | HSALG010569.2 | XM_011156821.3, NM_001169697.3 | Abdominal fat body | LOC105192591 |
| *Elovl* | fatty acid elongase AAEL008004, CG31522 | HSALG002373.2 | XM_011153792.3, NM_001259976.2 | Abdominal fat body | LOC105190836 |
| *Scd* | acyl-CoA Delta(11) desaturase | HSALG013065.2 | XM_011153192.2 | Abdominal fat body | LOC105190430 |
| *Fasn* | fatty acid synthase-like | HSALG002054.2 | XM_025305970.1, NM_001015405.4 | Abdominal fat body | LOC112590176 |
| *Egfr* | Epidermal Growth Factor Receptor | HSALG001375.2 | XM_011155066.3, NM_057410.4 | Abdominal fat body | LOC105191569 |
| *Vein* | EGFR ligand | HSALG004141.2 | XM_025297750.1, NM_001014565.2 | Abdominal fat body, Ovary | LOC105181399 |
| *Spitz* | EGFR ligand | HSALG008632.1 | XM_011146179.3, NM_001032110.2 | Abdominal fat body | LOC105186150 |

**Table S2.**
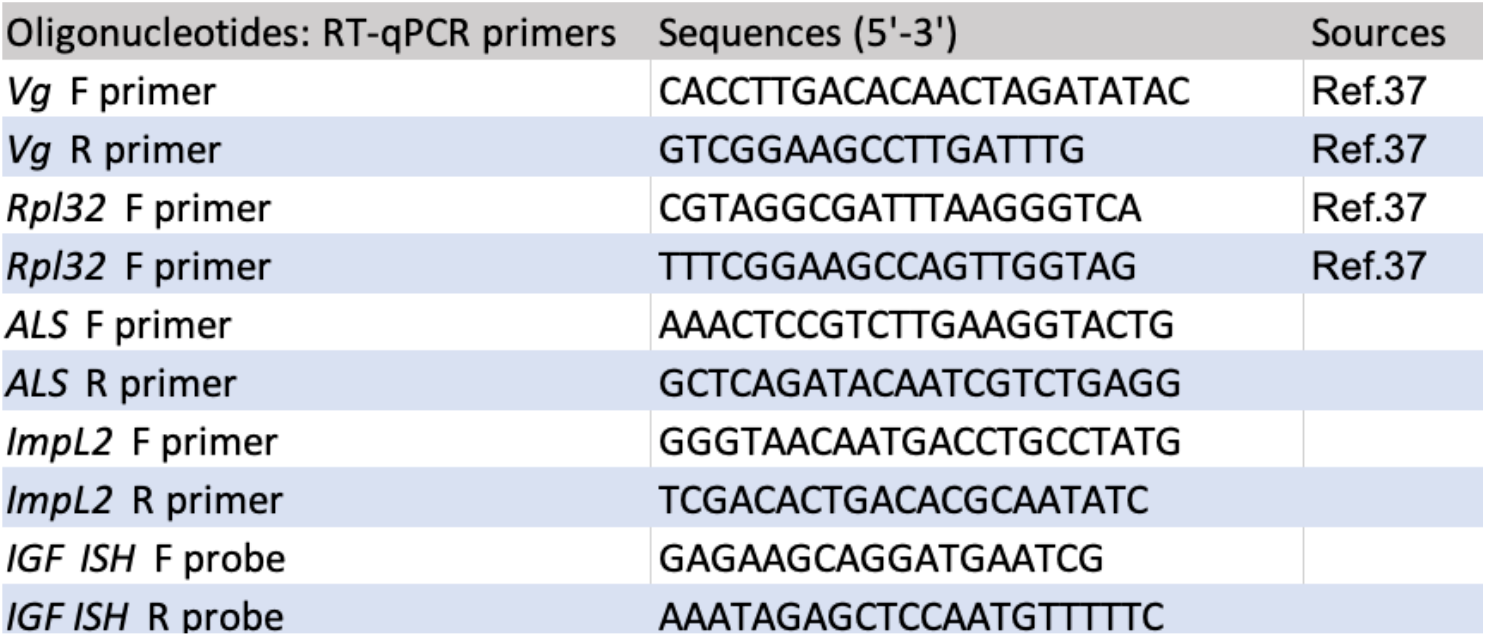
Oligo sequences and their sources (right column) used in this study.

## METHODS

### Regular maintenance of *Harpegnathos* ant colonies

*Harpegnathos saltator* colonies were initially transported from Jürgen Liebig’s laboratory at Arizona State University. Ants were housed in plastic boxes (Pioneer Plastics, Inc.) with a plaster floor (Darby Dental, #8491560) inside a 12:12 h light/dark cycle and temperature-controlled room (22-25°C). Two compartments in a box divide the nest area for reproductive females, young workers (nurses) and the broods from the foraging arena for old workers (foragers). Ants and their broods were fed with pre-stung (paralyzed) crickets three times a week.

### Caste transition (W to G) and reversion (G to R)

As shown in **Fig. S1A**., a queenless colony consisting of thirty of workers (∼2-4 weeks post emergence) were setup for inducing a gamergate transition for three months. Each individual was uniquely marked on thorax with Uni-paint markers. Well-known behavioral traits in gamergates: antennal dueling and egg laying events, were observed for identifying a mature gamergate (*35, 58, 88*). For the reversion, the mature gamergate (G), derived from 3 months of caste transition, was removed from the transition colony, individually isolated and constantly fed with small pre-stung crickets (1/4”) for 4 weeks. Egg laying events were observed in the isolated mature gamergates. Workers (W) from the same transition colony were placed in another mature colony with established reproductive females to prevent the gamergate transition after the gamergate removal and were used as a control worker in a lifespan experiment. The isolated gamergates were subsequently transferred into the mature colony where worker policing occurred. After two months, the reverted gamergates or revertants (R) fully exhibited worker-like behavior and stopped egg-laying (*36*). All tissues of workers, gamergates and revertants were harvested at the same chronological age (∼6 months old). Worker and gamergate samples were derived from the caste transition colony while revertants were independently from the reversion colony.

### Identification of insulin-like peptides (ILPs) in *Harpegnathos saltator*

The ILPs of *Drosophila* were used as queries to search against the genomes and/or transcriptomes of 36 representative hymenopteran insect species through the NCBI BLASTP and TBLASTN web interface (http://blast.ncbi.nlm.nih.gov/Blast.cgi). Multiple sequence alignment of all identified hymenopteran ILP homologs were created using the E-INS-i method of MAFFT v7.425 and filtered for gap-rich regions using TrimAl v1.4. A preliminary phylogenetic analysis was carried out using IQ-TREE v1.6.10 with 10 independent tree searches and branch supports were measured using the Ultrafast Bootstrap approach with 1000 replicates. The result clearly separated hymenopteran ILPs into two groups, namely Ins and IGF. Phylogenetic trees of Ins and IGF were then reconstructed, respectively, following the procedure described above.

### Transcriptome analysis and RT-qPCR

*Harpegnathos* tissues were dissected from single individuals and homogenized in TriPure (Sigma). Three different tissues from 6-month-old workers (W), gamergates (G) and revertants (R) were collected including the central brain without optic lobes, whole ovary and abdominal fat body. Total RNA was purified by Phenol/Chloroform extraction. RNA was reverse transcribed to cDNA using QuantiTect Reverse Transcription Kit (Qiagen).

For library preparation, polyA+ RNA was isolated from total RNA using Dynabeads Oligo(dT) 25 (Thermo Fisher) beads. The 1^st^ strand was synthesized by Superscript III and random hexamers (Life Technologies). The 2^nd^ strand was synthesized with dUTP to generate strand asymmetry using DNA Pol I (NEB, M0209L) and E. coli DNA Ligase (Enzymatics, L6090L) (*103*). RNA-seq libraries were constructed using the protocol described in (*104*). Sequencing was performed on a HiSeq 2500 (Illumina) at NYULMC.

Reads were aligned with STAR (v.2.5.0b) (*105*) on the *Harpegnathos* genome version 8.5 (*106*). Read numbers in CPM (Counts Per Million) and differential expression analysis were calculated and performed using EdgeR (*107*). An adjusted p-value of less than 0.05 was considered significant. Gene identifiers in the study were reported in Table S1.

For qPCR, mRNA expression levels were quantified using LightCycler^®^ 480 SYBR Green I Master (Roche#04707516001) with a specific primer pair (See Table S2). Ribosomal protein (Rpl32) and GAPDH genes were used as normalization controls.

### In situ hybridization (ISH) and immunofluorescence (IF) analyses

Tissue dissection was performed under 1x Phosphate Buffered Saline (PBS) supplemented with Protease and Phosphatase inhibitor cocktail (Thermo Scientific, #78443) on a clean, clear rubber pad using two fine forceps (Fine Science Tools, Dumont #5) and scissor (Fine Science Tools, #1500408). Dissected tissues were fixed using 4% Paraformaldehyde (PFA) (Electron Microscopy Sciences, #15710) diluted in 1x PBS with 0.1% Tween20 (0.1% PBST) at room temperature (RT) for 20 min on a shaker. Fixed tissues were washed three times with 0.1% PBST for 15 min each on a shaker.

#### ISH

Tissues were gradually dehydrated by incubating in serially diluted methanol at 25%, 50% and 90% at RT for 10 min each on a shaker. Tissues were rehydrated by performing a graded rehydration using 100%, 90%, 50%, and 25% methanol at RT for 10 min each. Tissue was then washed in 0.1% PBST twice for five min at RT while shaking. Post-fixation was performed by incubating in 4% PFA in 1x PBS for 20 min at RT on a shaker. Tissue was washed with PBST twice for five minutes while shaking. Tissue was permeabilized by incubating with Proteinase K (10 ug/ml) for two min at RT. Following incubation, the tissue was washed with 0.1% PBST for five min at RT while nutating. Another post fixation step was performed by incubating in 4% PFA in 1x PBS for 20 min at RT on a nutator. Tissue was subsequently washed with 0.1% PBST two times for 5 min at RT. The PBST was removed and replaced with pre-warmed hybridization buffer (50% formamide, 5x saline-sodium citrate (SSC), 5x Denhardt’s solution, 250 ug/ml yeast tRNA, 500 ug/ml herring sperm DNA, 50 ug/ml Heparin, 2.5 mM EDTA, 0.1% Tween20, and 0.25% CHAPS) at 55° C for approximately two hours. Digoxigenin (DIG) labeled probe was synthesized using the DIG RNA labeling kit (SP6/T7) (Roche, #11175025910). Denatured DIG-labeled probe was hybridized at 55° C overnight. Hybridized tissue was then washed twice with warm hybridization solution for 40 min at 55° C and washed four times with 0.1% PBST for 10 min at RT. Detection of DIG-labeled probes was performed by incubating with anti-DIG antibody (1:2000, Roche #11093274910) in 0.1% PBST overnight at 4° C. Tissue was washed three times with 0.1% PBST and washed twice with AP buffer (100 mM NaCl, 50 mM MgCl_2_, 100 mM Tris, pH 9.5, and 0.1% Tween20) for 10 min each at RT. Colorimetric detection was performed by incubating with NBT/BCIP solution (Roche, # 11697471001) for 10 min. The colorimetric reaction was halted by washing with 0.1% PBST two times for five min at RT while shaking.

#### IF

Fixed tissue was placed in custom antibodies of Ins, IGF, ALS, Imp-L2 and FOXO or pre-immune serum (1:100) in 0.1% PBSTx (TritonX-100) with 5% normal serum (Fig. S6C). Tissue was incubated overnight at 4° C on a nutator. Subsequent removal of primary antibody was performed and tissue was washed four times for five min at RT in 0.1% PBSTx. Tissue was probed with anti-rabbit, -rat and –guinea pig secondary antibodies at RT for 1 hour on a nutator (1:500, ThermoFisher, #A21206, A21470 and A11073, respectively) and washed four times for five min at RT in 0.1% PBSTx. Counterstaining was performed by incubating with 2 μg/ml DAPI and 1:100 Alexa Fluor™ 488 Phalloidin (ThermoFisher, #A12379) diluted in 0.1% PBSTx for 20 min at RT in the dark. Tissue was rinsed 5 times with 0.1% PBSTx, mounted in Focusclear and imaged using a Zeiss 880 Confocal microscope at NYULH.

### Western analysis

Anti-*Drosophila* phospho-AKT (Ser505) and anti-AKT antibodies have been used in *Drosophila* and other insects, such as the silkworm *Bombyx mori* (*108*) and the planthopper *Nilaparvata lugens* (*109*).

Dissected tissues of foraging workers and mature gamergates were homogenized with lysis buffer (50 mM Tris-HCl, pH 7.8, 150 mM NaCl, and 1% Nonidet P-40) plus Halt protease/phosphatase inhibitor cocktail (78442). Approximately 10 μg of tissue lysate was loaded onto 8% Bis-Tris gels. PVDF membranes were used and blocked in TBST buffer (150 mM NaCl, 20 mM Tris-HCl, pH 7.5, and 0.05% v/v Tween20) containing 3% milk (Carnation powdered milk) for 1 hour at RT and probed overnight at 4° C with primary rabbit antibodies (anti-phospho-AKT #4054S, anti-phospho MAPK #9101S, two anti-total-AKT #9272S and #4691S, and anti-total-MAPK#4695S (Cell Signaling Technology, 1:2000), guinea pig anti-ImpL2 (1:1000) and rat anti-ALS (1:1000), diluted in TBST with 5% BSA and 0.04% sodium azide (NaN_3_). Anti-rabbit (Promega, #W401B), anti-guinea pig (JacksonImmuno, #106035003) and anti-rat IgG HRP conjugate secondary antibodies (JacksonImmuno, #715035150) were used at a 1:5000 dilution in TBST containing 5% BSA or 3% milk. West FEMTO Maximum Sensitivity Substrate (Thermo Fisher, #34094) was used for signal detection. Membrane-bound antibodies were removed using Western Blot Stripping buffer (ThermoFisher, # 21059). Band intensity was quantified using Fiji. Unpaired t test was performed using GraphPad Prism software.

### Treatment of ant tissues by synthetic ILP peptides

Chemically synthetic Ins peptides were prepared by Phoenix Pharmaceuticals Inc. Two peptides (A and B chains) were synthesized and 3 disulfide bonds were formed.

A Chain: ARGIYEECCLNACTYNELSTYCGPQQ;

B Chain: QSDGSYALKWSMNVPQRYCGRRLSNALQTVCTGVYNNMF;

Disulfide bonds: [CysA9-CysB19], [CysA22-CysB31], [CysA8-CysA13].

Abdominal fat body tissues of workers were dissected in serum-free Schneider’s medium (Invitrogen, #21720) supplemented with 1% streptomycin/penicillin. After 4 hours of starvation in serum-free medium, chemically synthetic Ins peptides were added to the media. After 30 min of treatment, media were discarded and protein lysates from tissues were subjected for Western blots. One-way ANOVA with Tukey’s multiple comparisons test were performed using GraphPad Prism software.

### Injections of Insulin peptide and MAPK inhibitor (U0126)

For the *in vivo* experiment, 40 workers, two-weeks post-pupal emergence, were individually labeled with distinct paint dots and housed in a queenless condition. We injected approximately 1 µl of 100 µM synthetic insulin peptide into the worker abdomen (an opening between tergites) two days after setting up the colonies. Controls were injected with water. Antennal dueling behavior were observed for 5 days post-dueling initiation. All individuals were sacrificed after 5 days of the dueling tournament for ovary scoring by counting the number of yolky oocytes. For the U0126 experiment, workers were injected with approximately 1 µl of 100 µM U0126 or 1% DMSO diluted with water. The ovary and abdominal fat body were collected six days post-injection for scoring ovary development and quantifying *vitellogenin* expression, respectively. Injections of both 100 µM U0126 and insulin peptide were performed separately. Half of the workers (20 workers) was first injected with either U0126 or 1% DMSO (control). Insulin was subsequently injected in all workers (40 workers) on the following day. Dueling activities were monitored for 5 days and ovary development was scored 5 days post-dueling tournament. Three independent colonies were performed. Animals that did not survive the injections were excluded.

### ALS and Imp-L2 protein production using baculovirus expression system

To purify ovarian ALS and ImpL-2 proteins, Flag/HIS-tagged ALS and Imp-L2 were independently cloned into a baculovirus expression vector, pFASTBac1 (Invitrogen) and expressed in Sf9 cells as described previously (*110*). Infected Sf9 cells were resuspended in BC150 buffer (25 mM HEPES-NaOH, pH 7.5, 1 mM EDTA, 150 mM NaCl, 10% glycerol, 1 mM DTT, and 0.1% NP-40) with protease inhibitors (1 mM phenylmethlysulfonyl fluoride [PMSF], 0.1 mM benzamidine, 1.25 mg/mL leupeptin, and 0.625 mg/mL pepstatin A) and phosphatase inhibitors (20 mM NaF and 1 mM Na3VO4). Cells were lysed by sonication (Fisher Sonic Dismembrator model 100) and purified through FLAG-M2 agarose beads (Sigma).

### Lipid and Sugar Quantifications from the Tissues and Hemolymph

Ants were rinsed with ultrapure water and anesthetized on ice. Extra liquid was removed. An abdomen and all legs were clipped off from the whole body. The abdomen obtained was dissected for the abdominal fat body and ovary under ice cold 1x PBS. The dissected tissues were homogenized in ice cold 1xPBS using an insulin needle (BD, #328291) or a pestle. The head and thorax were placed in a 0.5 ml tube with a small hole in the bottom inside a 1.5 ml collection tube. Hemolymph was collected after centrifugation at 3,000xg for 1 minute at 4° C. Approximately 1 μl of clear hemolymph was subjected to each lipid and sugar measurement. Triglyceride quantification was performed using Infinity Triglycerides reagent (Thermofisher, #TR-22421), as described in the manufacturer’s instructions and (*111*). For sugar (trehalose and glucose) measurements, the lysates were diluted in trehalose buffer (5 mM Tris pH 6.6, 137 mM NaCl, and 2.7 mM KCl). The samples obtained were incubated with porcine trehalase (Sigma, #T8778) at 37° C overnight. Measurements of trehalose and glucose levels were performed using the Infinity Glucose Hexokinase reagent (Thermofisher, #TR-15421), as described in the manufacturer’s instructions and (*111*).

### Quantification and Statistical Analysis

Statistical analysis of transcriptome data was performed using EdgeR. Other statistical analyses, including unpaired t test, one-way ANOVA with Tukey’s multiple comparisons test, and Log-rank Mantel Cox test, as indicated in Methods, were performed using GraphPad Prism software. The value of n, mean ± SEM, and p value are reported in Results, Figures, and Figure Legends. Statistical significance is defined by p<0.05 (*), p<0.01 (**), or p<0.001 (***).

## Notes

### Competing Interest Statement

The authors have declared no competing interest.

## References

1. N. L. Jenkins, G. McColl, G. J. Lithgow, Fitness cost of extended lifespan in Caenorhabditis elegans. Proc Biol Sci 271, 2523–2526 (2004).

2. M. Ezcurra et al., C. elegans Eats Its Own Intestine to Make Yolk Leading to Multiple Senescent Pathologies. Curr Biol 28, 2544–2556 e2545 (2018).

3. A. W. McCracken, G. Adams, L. Hartshorne, M. Tatar, M. J. P. Simons, The hidden costs of dietary restriction: Implications for its evolutionary and mechanistic origins. Sci Adv 6, eaay3047 (2020).

4. S. C. Johnson, P. S. Rabinovitch, M. Kaeberlein, mTOR is a key modulator of ageing and age-related disease. Nature 493, 338–345 (2013).

5. E. H. Chen, Q. L. Hou, D. D. Wei, H. B. Jiang, J. J. Wang, Phenotypic plasticity, trade-offs and gene expression changes accompanying dietary restriction and switches in Bactrocera dorsalis (Hendel) (Diptera: Tephritidae). Sci Rep 7, 1988 (2017).

6. G. E. Blomquist, Trade-off between age of first reproduction and survival in a female primate. Biology letters 5, 339–342 (2009).

7. T. Flatt, M. P. Tu, M. Tatar, Hormonal pleiotropy and the juvenile hormone regulation of Drosophila development and life history. BioEssays : news and reviews in molecular, cellular and developmental biology 27, 999–1010 (2005).

8. M. Tatar, The plate half-full: status of research on the mechanisms of dietary restriction in Drosophila melanogaster. Experimental gerontology 46, 363–368 (2011).

9. R. G. J. Westendorp, T. B. L. Kirkwood, Human longevity at the cost of reproductive success. Nature 396, 743–746 (1998).

10. M. Tatar et al., A mutant Drosophila insulin receptor homolog that extends life-span and impairs neuroendocrine function. Science 292, 107–110 (2001).

11. K. D. Kimura, H. A. Tissenbaum, Y. Liu, G. Ruvkun, daf-2, an insulin receptor-like gene that regulates longevity and diapause in Caenorhabditis elegans. Science 277, 942–946 (1997).

12. L. Fontana, L. Partridge, V. D. Longo, Extending healthy life span--from yeast to humans. Science 328, 321–326 (2010).

13. M. Bluher, B. B. Kahn, C. R. Kahn, Extended longevity in mice lacking the insulin receptor in adipose tissue. Science 299, 572–574 (2003).

14. M. Tatar, A. Bartke, A. Antebi, The endocrine regulation of aging by insulin-like signals. Science 299, 1346–1351 (2003).

15. L. Partridge, N. Alic, I. Bjedov, M. D. Piper, Ageing in Drosophila: the role of the insulin/Igf and TOR signalling network. Experimental gerontology 46, 376–381 (2011).

16. S. J. Broughton et al., Longer lifespan, altered metabolism, and stress resistance in Drosophila from ablation of cells making insulin-like ligands. Proc Natl Acad Sci U S A 102, 3105–3110 (2005).

17. E. L. Arrese, J. L. Soulages, Insect fat body: energy, metabolism, and regulation. Annu Rev Entomol 55, 207–225 (2010).

18. E. Gutierrez, D. Wiggins, B. Fielding, A. P. Gould, Specialized hepatocyte-like cells regulate Drosophila lipid metabolism. Nature 445, 275–280 (2007).

19. M. Bluher, Fat tissue and long life. Obesity facts 1, 176–182 (2008).

20. C. Geminard, E. J. Rulifson, P. Leopold, Remote control of insulin secretion by fat cells in Drosophila. Cell metabolism 10, 199–207 (2009).

21. M. E. Giannakou et al., Long-lived Drosophila with overexpressed dFOXO in adult fat body. Science 305, 361 (2004).

22. L. LaFever, D. Drummond-Barbosa, Direct control of germline stem cell division and cyst growth by neural insulin in Drosophila. Science 309, 1071–1073 (2005).

23. D. S. Hwangbo, B. Gershman, M. P. Tu, M. Palmer, M. Tatar, Drosophila dFOXO controls lifespan and regulates insulin signalling in brain and fat body. Nature 429, 562–566 (2004).

24. S. Gronke, D. F. Clarke, S. Broughton, T. D. Andrews, L. Partridge, Molecular evolution and functional characterization of Drosophila insulin-like peptides. PLoS Genet 6, e1000857 (2010).

25. C. M. Sgro, L. Partridge, A delayed wave of death from reproduction in Drosophila. Science 286, 2521–2524 (1999).

26. H. Hsin, C. Kenyon, Signals from the reproductive system regulate the lifespan of C. elegans. Nature 399, 362–366 (1999).

27. N. Arantes-Oliveira, J. Apfeld, A. Dillin, C. Kenyon, Regulation of life-span by germ-line stem cells in Caenorhabditis elegans. Science 295, 502–505 (2002).

28. L. Keller, M. Genoud, Extraordinary lifespans in ants: a test of evolutionary theories of ageing. Nature 389, 958–960 (1997).

29. M. Ghaninia et al., Chemosensory sensitivity reflects reproductive status in the ant Harpegnathos saltator. Sci Rep 7, 3732 (2017).

30. B. Holldobler, E. O. Wilson, The Ants. (Belknap Press: An Imprint of Harvard University Press, 1990).

31. P. Calabi, S. D. Porter, Worker longevity in the fire ant Solenopsis invicta: Ergonomic considerations of correlations between temperature, size and metabolic rates. Journal of Insect Physiology 35, 643–649 (1989).

32. H. Yan et al., Eusocial insects as emerging models for behavioural epigenetics. Nat Rev Genet 15, 677–688 (2014).

33. C. Opachaloemphan, H. Yan, A. Leibholz, C. Desplan, D. Reinberg, Recent Advances in Behavioral (Epi)Genetics in Eusocial Insects. Annual review of genetics 52, 489–510 (2018).

34. J. Heinze, B. Hölldobler, C. Peeters, Conflict and Cooperation in Ant Societies. Naturwissenschaften 81, 489–497 (1994).

35. C. Opachaloemphan et al., Early behavioral and molecular events leading to caste switching in the ant Harpegnathos. Genes & development 35, 410–424 (2021).

36. C. A. Penick et al., Reversible plasticity in brain size, behaviour and physiology characterizes caste transitions in a socially flexible ant (Harpegnathos saltator). Proc Biol Sci 288, 20210141 (2021).

37. J. Gospocic et al., The Neuropeptide Corazonin Controls Social Behavior and Caste Identity in Ants. Cell 170, 748–759 e712 (2017).

38. D. J. Burks et al., IRS-2 pathways integrate female reproduction and energy homeostasis. Nature 407, 377–382 (2000).

39. W. Brogiolo et al., An evolutionarily conserved function of the Drosophila insulin receptor and insulin-like peptides in growth control. Current biology : CB 11, 213–221 (2001).

40. T. Fujii et al., Sex pheromone desaturase functioning in a primitive Ostrinia moth is cryptically conserved in congeners’ genomes. Proc Natl Acad Sci U S A 108, 7102–7106 (2011).

41. T. Chertemps et al., A female-biased expressed elongase involved in long-chain hydrocarbon biosynthesis and courtship behavior in Drosophila melanogaster. Proc Natl Acad Sci U S A 104, 4273–4278 (2007).

42. J. M. Lassance et al., Allelic variation in a fatty-acyl reductase gene causes divergence in moth sex pheromones. Nature 466, 486–489 (2010).

43. J. Liebig, C. Peeters, N. J. Oldham, C. Markstadter, B. Hölldobler, Are variations in cuticular hydrocarbons of queens and workers a reliable signal of fertility in the ant Harpegnathos saltator? Proc Natl Acad Sci U S A 97, 4124–4131 (2000).

44. A. S. Raikhel, T. S. Dhadialla, Accumulation of yolk proteins in insect oocytes. Annu Rev Entomol 37, 217–251 (1992).

45. H. Yan, J. Liebig, Genetic basis of chemical communication in eusocial insects. Genes & development 35, 470–482 (2021).

46. J. F. Ferveur et al., Genetic feminization of pheromones and its behavioral consequences in Drosophila males. Science 276, 1555–1558 (1997).

47. V. Chandra et al., Social regulation of insulin signaling and the evolution of eusociality in ants. Science 361, 398–402 (2018).

48. R. S. Garofalo, Genetic analysis of insulin signaling in Drosophila. Trends in Endocrinology & Metabolism 13, 156–162 (2002).

49. V. M. Fernandes, Z. Chen, A. M. Rossi, J. Zipfel, C. Desplan, Glia relay differentiation cues to coordinate neuronal development in Drosophila. Science 357, 886–891 (2017).

50. C. Slack et al., The Ras-Erk-ETS-Signaling Pathway Is a Drug Target for Longevity. Cell 162, 72–83 (2015).

51. C. Slack, M. E. Giannakou, A. Foley, M. Goss, L. Partridge, dFOXO-independent effects of reduced insulin-like signaling in Drosophila. Aging Cell 10, 735–748 (2011).

52. K. E. Brown, M. Kerr, M. Freeman, The EGFR ligands Spitz and Keren act cooperatively in the Drosophila eye. Dev Biol 307, 105–113 (2007).

53. B. Schnepp, G. Grumbling, T. Donaldson, A. Simcox, Vein is a novel component in the Drosophila epidermal growth factor receptor pathway with similarity to the neuregulins. Genes Dev 10, 2302–2313 (1996).

54. P. Decio, A. S. Vieira, N. B. Dias, M. S. Palma, O. C. Bueno, The Postpharyngeal Gland: Specialized Organ for Lipid Nutrition in Leaf-Cutting Ants. PLoS One 11, e0154891 (2016).

55. D. Eelen, L. Børgesen, J. Billen, Functional morphology of the postpharyngeal gland of queens and workers of the ant Monomorium pharaonis (L.). Acta Zoologica 87, 101–111 (2006).

56. S. S. Lee, S. Kennedy, A. C. Tolonen, G. Ruvkun, DAF-16 target genes that control C. elegans life-span and metabolism. Science 300, 644–647 (2003).

57. A. A. Parkhitko et al., Downregulation of the tyrosine degradation pathway extends Drosophila lifespan. Elife 9, (2020).

58. C. A. Penick, C. S. Brent, K. Dolezal, J. Liebig, Neurohormonal changes associated with ritualized combat and the formation of a reproductive hierarchy in the ant Harpegnathos saltator. J Exp Biol 217, 1496–1503 (2014).

59. H. J. Hsu, D. Drummond-Barbosa, Insulin levels control female germline stem cell maintenance via the niche in Drosophila. Proc Natl Acad Sci U S A 106, 1117–1121 (2009).

60. S. Luo, G. A. Kleemann, J. M. Ashraf, W. M. Shaw, C. T. Murphy, TGF-beta and insulin signaling regulate reproductive aging via oocyte and germline quality maintenance. Cell 143, 299–312 (2010).

61. D. Michaelson, D. Z. Korta, Y. Capua, E. J. Hubbard, Insulin signaling promotes germline proliferation in C. elegans. Development 137, 671–680 (2010).

62. N. Arquier et al., Drosophila ALS regulates growth and metabolism through functional interaction with insulin-like peptides. Cell metabolism 7, 333–338 (2008).

63. B. Honegger et al., Imp-L2, a putative homolog of vertebrate IGF-binding protein 7, counteracts insulin signaling in Drosophila and is essential for starvation resistance. J Biol 7, 10 (2008).

64. N. Okamoto et al., A secreted decoy of InR antagonizes insulin/IGF signaling to restrict body growth in Drosophila. Genes Dev 27, 87–97 (2013).

65. W. Sajid et al., Structural and biological properties of the Drosophila insulin-like peptide 5 show evolutionary conservation. J Biol Chem 286, 661–673 (2011).

66. A. Sloth Andersen, P. Hertz Hansen, L. Schaffer, C. Kristensen, A new secreted insect protein belonging to the immunoglobulin superfamily binds insulin and related peptides and inhibits their activities. J Biol Chem 275, 16948–16953 (2000).

67. Y. Yamanaka, E. M. Wilson, R. G. Rosenfeld, Y. Oh, Inhibition of insulin receptor activation by insulin-like growth factor binding proteins. J Biol Chem 272, 30729–30734 (1997).

68. I. Ueki et al., Inactivation of the acid labile subunit gene in mice results in mild retardation of postnatal growth despite profound disruptions in the circulating insulin-like growth factor system. Proc Natl Acad Sci U S A 97, 6868–6873 (2000).

69. H. M. Domene et al., Phenotypic effects of null and haploinsufficiency of acid-labile subunit in a family with two novel IGFALS gene mutations. J Clin Endocrinol Metab 92, 4444–4450 (2007).

70. J. Dai, R. C. Baxter, Regulation in vivo of the acid-labile subunit of the rat serum insulin-like growth factor-binding protein complex. Endocrinology 135, 2335–2341 (1994).

71. S. A. Wandji, J. E. Gadsby, F. A. Simmen, J. A. Barber, J. M. Hammond, Porcine ovarian cells express messenger ribonucleic acids for the acid-labile subunit and insulin-like growth factor binding protein-3 during follicular and luteal phases of the estrous cycle. Endocrinology 141, 2638–2647 (2000).

72. E. Chin, J. Zhou, J. Dai, R. C. Baxter, C. A. Bondy, Cellular localization and regulation of gene expression for components of the insulin-like growth factor ternary binding protein complex. Endocrinology 134, 2498–2504 (1994).

73. L. Manning et al., A hormonal cue promotes timely follicle cell migration by modulating transcription profiles. Mech Dev 148, 56–68 (2017).

74. V. Evdokimova et al., IGFBP7 binds to the IGF-1 receptor and blocks its activation by insulin-like growth factors. Sci Signal 5, ra92 (2012).

75. J. Liebig, H.-J. Poethke, Queen lifespan and colony longevity in the ant Harpegnathos saltator. Ecological Entomology 29, 203–207 (2004).

76. K. D. Baker, C. S. Thummel, Diabetic larvae and obese flies-emerging studies of metabolism in Drosophila. Cell Metab 6, 257–266 (2007).

77. A. R. Saltiel, C. R. Kahn, Insulin signalling and the regulation of glucose and lipid metabolism. Nature 414, 799–806 (2001).

78. J. A. Engelman, J. Luo, L. C. Cantley, The evolution of phosphatidylinositol 3-kinases as regulators of growth and metabolism. Nat Rev Genet 7, 606–619 (2006).

79. R. L. Elstrom et al., Akt stimulates aerobic glycolysis in cancer cells. Cancer Res 64, 3892–3899 (2004).

80. H. Huang, D. J. Tindall, Dynamic FoxO transcription factors. J Cell Sci 120, 2479–2487 (2007).

81. D. H. Kim et al., FoxO6 integrates insulin signaling with MTP for regulating VLDL production in the liver. Endocrinology 155, 1255–1267 (2014).

82. A. Kamagate et al., FoxO1 mediates insulin-dependent regulation of hepatic VLDL production in mice. J Clin Invest 118, 2347–2364 (2008).

83. L. Shi, B. P. Tu, Acetyl-CoA and the regulation of metabolism: mechanisms and consequences. Curr Opin Cell Biol 33, 125–131 (2015).

84. F. Pietrocola, L. Galluzzi, J. M. Bravo-San Pedro, F. Madeo, G. Kroemer, Acetyl coenzyme A: a central metabolite and second messenger. Cell Metab 21, 805–821 (2015).

85. W. Gronenberg, J. Liebig, Smaller brains and optic lobes in reproductive workers of the ant Harpegnathos. Naturwissenschaften 86, 343–345 (1999).

86. A. R. Armstrong, D. Drummond-Barbosa, Insulin signaling acts in adult adipocytes via GSK-3beta and independently of FOXO to control Drosophila female germline stem cell numbers. Dev Biol 440, 31–39 (2018).

87. Y. Zhang et al., Irisin stimulates browning of white adipocytes through mitogen-activated protein kinase p38 MAP kinase and ERK MAP kinase signaling. Diabetes 63, 514–525 (2014).

88. C. A. Penick, J. Liebig, C. S. Brent, Reproduction, dominance, and caste: endocrine profiles of queens and workers of the ant Harpegnathos saltator. Journal of comparative physiology. A, Neuroethology, sensory, neural, and behavioral physiology 197, 1063–1071 (2011).

89. L. Zipper, D. Jassmann, S. Burgmer, B. Gorlich, T. Reiff, Ecdysone steroid hormone remote controls intestinal stem cell fate decisions via the PPARgamma-homolog Eip75B in Drosophila. Elife 9, (2020).

90. S. M. H. Ahmed et al., Fitness trade-offs incurred by ovary-to-gut steroid signalling in Drosophila. Nature 584, 415–419 (2020).

91. J. Cruz, D. Martin, X. Franch-Marro, Egfr Signaling Is a Major Regulator of Ecdysone Biosynthesis in the Drosophila Prothoracic Gland. Curr Biol 30, 1547–1554 e1544 (2020).

92. D. L. Osterbur, D. K. Fristrom, J. E. Natzle, S. J. Tojo, J. W. Fristrom, Genes expressed during imaginal discs morphogenesis: IMP-L2, a gene expressed during imaginal disc and imaginal histoblast morphogenesis. Developmental Biology 129, 439–448 (1988).

93. M. Ghaninia et al., Antennal Olfactory Physiology and Behavior of Males of the Ponerine Ant Harpegnathos saltator. J Chem Ecol 44, 999–1007 (2018).

94. V. Dietemann, C. Peeters, J. Liebig, V. Thivet, B. Holldobler, Cuticular hydrocarbons mediate discrimination of reproductives and nonreproductives in the ant Myrmecia gulosa. Proc Natl Acad Sci U S A 100, 10341–10346 (2003).

95. S. Ahmed et al., Identification of membrane-bound serine proteinase matriptase as processing enzyme of insulin-like growth factor binding protein-related protein-1 (IGFBP-rP1/angiomodulin/mac25). FEBS J 273, 615–627 (2006).

96. L. Drees et al., Conserved function of the matriptase-prostasin proteolytic cascade during epithelial morphogenesis. PLoS Genet 15, e1007882 (2019).

97. A. Figueroa-Clarevega, D. Bilder, Malignant Drosophila tumors interrupt insulin signaling to induce cachexia-like wasting. Dev Cell 33, 47–55 (2015).

98. Y. Kwon et al., Systemic organ wasting induced by localized expression of the secreted insulin/IGF antagonist ImpL2. Dev Cell 33, 36–46 (2015).

99. M. S. Dionne, L. N. Pham, M. Shirasu-Hiza, D. S. Schneider, Akt and FOXO dysregulation contribute to infection-induced wasting in Drosophila. Curr Biol 16, 1977–1985 (2006).

100. J. Lee, K. G. Ng, K. M. Dombek, D. S. Eom, Y. V. Kwon, Tumors overcome the action of the wasting factor ImpL2 by locally elevating Wnt/Wingless. Proc Natl Acad Sci U S A 118, (2021).

101. M. V. Blagosklonny, Aging and immortality: quasi-programmed senescence and its pharmacologic inhibition. Cell Cycle 5, 2087–2102 (2006).

102. D. Gems, L. Partridge, Genetics of longevity in model organisms: debates and paradigm shifts. Annu Rev Physiol 75, 621–644 (2013).

103. D. Parkhomchuk et al., Transcriptome analysis by strand-specific sequencing of complementary DNA. Nucleic Acids Res 37, e123 (2009).

104. V. Narendra et al., CTCF establishes discrete functional chromatin domains at the Hox clusters during differentiation. Science 347, 1017–1021 (2015).

105. A. Dobin et al., STAR: ultrafast universal RNA-seq aligner. Bioinformatics 29, 15–21 (2013).

106. E. J. Shields, L. Sheng, A. K. Weiner, B. A. Garcia, R. Bonasio, High-Quality Genome Assemblies Reveal Long Non-coding RNAs Expressed in Ant Brains. Cell reports 23, 3078–3090 (2018).

107. M. I. Love, W. Huber, S. Anders, Moderated estimation of fold change and dispersion for RNA-seq data with DESeq2. Genome Biol 15, 550 (2014).

108. Y. Li et al., DNA synthesis during endomitosis is stimulated by insulin via the PI3K/Akt and TOR signaling pathways in the silk gland cells of Bombyx mori. International journal of molecular sciences 16, 6266–6280 (2015).

109. H. J. Xu et al., Two insulin receptors determine alternative wing morphs in planthoppers. Nature 519, 464–467 (2015).

110. C. H. Lee et al., Automethylation of PRC2 promotes H3K27 methylation and is impaired in H3K27M pediatric glioma. Genes Dev 33, 1428–1440 (2019).

111. J. M. Tennessen, W. E. Barry, J. Cox, C. S. Thummel, Methods for studying metabolism in Drosophila. Methods 68, 105–115 (2014).

